# EEG and pupillometric signatures of working memory overload

**DOI:** 10.1101/2022.11.21.517347

**Authors:** Alexandra I. Kosachenko, Dauren Kasanov, Alexander I. Kotyusov, Yuri G. Pavlov

## Abstract

Understanding the physiological correlates of cognitive overload has implications for gauging the limits of human cognition, for developing novel methods to define cognitive overload, and for mitigating the negative outcomes associated with overload. Most previous psychophysiological studies manipulated verbal working memory load in a narrow range (an average load of 5 items). It is unclear, however, how the nervous system responds to working memory load exceeding typical capacity limits. The objective of the current study was to characterize the central and autonomic nervous system changes associated with memory overload, by means of combined recording of EEG and pupillometry. Eighty-six participants were presented with a digit span task involving the serial auditory presentation of items. Each trial consisted of sequences of either 5, 9, or 13 digits, each separated by 2 seconds. Both theta activity and pupil size, after the initial rise, expressed a pattern of a short plateau and a decrease with reaching the state of memory overload, indicating that pupil size and theta possibly have similar neural mechanisms. Based on the described above triphasic pattern of pupil size temporal dynamics, we concluded that cognitive overload causes physiological systems to reset, and to release effort. Although memory capacity limits were exceeded and effort was released (as indicated by pupil dilation), alpha continued to decrease with increasing memory load. These results suggest that associating alpha with the focus of attention and distractor suppression is not warranted.

## Introduction

The human pupil dilates during effortful cognitive activities such as those that require increasing of deployed attentional control resources (Chen & Epps, 2014; Kursawe & Zimmer, 2015; Laeng et al., 2011; Lisi et al., 2015; Marquart & de Winter, 2015; Meghanathan et al., 2014). For example, maintaining information in working memory (WM) is accompanied by subjectively noticeable cognitive effort, and an increase in memory set size and cognitive effort is associated with pupil dilation during WM tasks (Heitz et al., 2007; Kahneman & Beatty, 1966; Karatekin et al., 2004; Klingner et al., 2011; Unsworth & Robison, 2015). Most previous pupillometric studies of cognitive load, employing the digit span task, showed a linear relationship between the number of presented memory items and pupil size (Karatekin, 2004; Klingner et al., 2011; Piquado et al., 2010). However, the majority used modest set sizes (4-6 items), while only a few studies used set sizes that exceeded the memory capacity, leading to cognitive overload.

How exactly does the pupil respond to cognitive overload? Two conflicting patterns of results have emerged over the years (Figure 1). Several studies have found an inverted U-shaped pattern where increasing memory load up to the capacity limit is associated with increasing pupil dilation, while further memory overload is associated with pupil constriction (Granholm et al., 1996; Johnson et al., 2014; Poock, 1973). By contrast, other studies have found a simpler, bi-phasic pattern whereby pupil size increases with load and reaches a stable plateau at the capacity limit (Cabestrero et al., 2009; Peavler, 1974). Given the association between pupil size and arousal, the exact shape of this pattern is indicative of how people respond to cognitive (over-)load: while an inverted U-shaped pattern would imply that cognitive and physiological effort are released, a prolonged plateau would imply that they are maintained even when task demands exceed the capacity limit.

**Figure 1.**
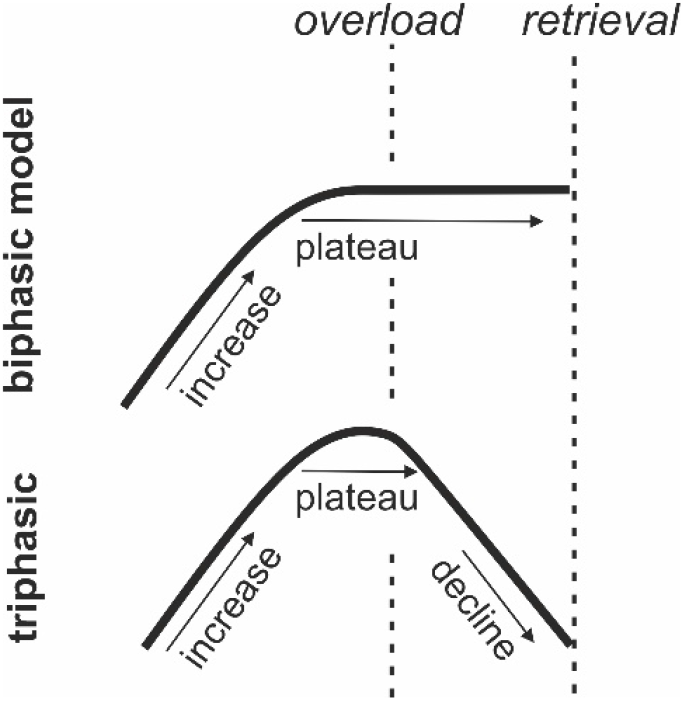
Two models of cognitive overload.

In the current study, we replicated the original Peavler’s (1974) experiment presenting participants with sequences of 5, 9, or 13 digits. We aimed to determine whether at the end of the retention interval after presentation of the 13th digit, pupil size would remain at the peak level or return to baseline. By using the simple span task with serial auditory presentation, we were able to monitor WM loading at a single item level. In deviation from the original protocol, we significantly increased the number of trials and expanded the sample size to increase signal-to-noise ratio.

The larger number of trials and the expanded sample size also allowed us to conduct an exploratory analysis of neural correlates of the cognitive overload state. To this end, we recorded electroencephalogram data concurrently with pupil diameter data. The application of EEG in a task with sequential presentation of memory items and sufficient duration between stimulus onsets (i.e., stimulus onset asynchrony; SOA) opened opportunities for exploring memory processes such as encoding and maintenance on a fine temporal scale. Our exploratory analysis of EEG data aimed to answer the following research questions:

### 1. Load and overload effects on alpha and theta activity

Individual WM performance is supported by two systems: the ability to control attention – also known as central executive, cognitive control, executive control, or executive components of WM (Baddeley, 2012; Bastian et al., 2020; Engle & Kane, 2004), and short-term storage with limited capacity. The short-term storage capacity refers to the amount of information an individual can maintain in the focus of attention or in a prioritized state at one time (Cowan, 2001). Both systems are necessary for successful task performance (Shipstead et al., 2014), but little is known about how they respond to WM overload.

Executive components of WM are frequently associated with frontal midline theta activity (Berger et al., 2014; Griesmayr et al., 2010; Pavlov & Kotchoubey, 2017; Sauseng et al., 2010). An increase in WM load is frequently accompanied by a linear increase in the power of theta activity (Pavlov & Kotchoubey, 2022), presumably, due to an increasing demand on executive control networks in the brain. It remains unclear whether demands on the executive control that exceed WM capacity prompt a breakdown of the network, or alternatively, prompt activity that remains constant at the level of individual memory capacity. These alternative outcomes may be reflected in the same tri- and biphasic models described above for pupil size, respectively. To date, however, these predictions have not been systematically investigated.

Moreover, WM retention has an effect on alpha power, although the direction of this effect is inconsistent across studies (Pavlov & Kotchoubey, 2022). Several studies have found suppression of alpha power during the retention interval (Erickson et al., 2019; Foster et al., 2016; Fukuda et al., 2015). Given the association between the alpha rhythm and cortical inhibition, these finding are consistent with greater activation of task-relevant cortical areas with higher WM demands. By contrast, several studies have found the opposite: an increase in alpha power during retention, which has been interpreted as greater inhibition of task-irrelevant information or of task-irrelevant cortical areas (Jensen & Mazaheri, 2010). Addressing this discrepancy, and characterizing the unexplored effect of WM overload, is essential for understanding the role of alpha activity in WM. Specifically, if alpha is a direct representation of the focus of attention or short-term storage (characterized by severely limited capacity) then we should not expect additional alpha suppression at very high levels of WM load (e.g., 10-13 digits) as compared to moderate levels of WM load (e.g., 6-9 digits). With that, protection from distractors role of alpha activity (and expected increase in alpha power) should express itself during retention interval (especially after the last digit in the sequence) to protect the memory trace from potentially distracting sensory input. Similarly, alpha enhancement should be more pronounced in individuals who actively suppress encoding of later digits in the sequence as their memory strategy. Finding an opposite pattern, would require us to reevaluate the functional role of alpha oscillations in WM.

To address these questions, we aimed to explore the relationship between cognitive load and frontal midline theta and posterior alpha power (a) within a typical load range (i.e., memorizing up to 7-9 digits) and (b) during cognitive overload (i.e., memorizing more than 9 digits).

### 2. Memory vs perception

Brain processes accompanying WM retention are typically confounded with sensory, memory-unrelated, processes, but it is uncommon to include a purely perceptual task as a control condition in a WM study. The only available EEG study that sequentially presented WM items (which would allow assessing oscillatory brain activity corresponding to encoding and retention of individual items) had a 1 second stimulus onset asynchrony (SOA) between items (Onton et al., 2005), which might be too short to reliably separate short-lived sensory from longer-lasting memory effects. To explore the oscillatory brain activity in the alpha and theta frequency bands during the task, we added a control condition with passive listening and employed a longer SOA of 2 seconds.

We assumed that the lack of a difference between control and memory conditions would suggest that the elicited oscillations are better explained by perceptual rather than memory processes.

### 3. Associations between pupil size, brain oscillations, and WM

It is possible that changes in oscillatory brain activity and pupil size can be explained by similar neurophysiological mechanisms. Pupil size has been proposed as an indirect measure of the locus coeruleus norepinephrine system (LC-NE) activity (Joshi et al., 2016; Murphy et al., 2014). Several studies reported a correlation between pupil dilation and alpha suppression during aversive learning and resting state, suggesting a contribution of LC-NE system in alpha suppression (Dahl et al., 2022; Sharon et al., 2021). Individual differences in WM capacity and attention control have been hypothesized to be driven by the differences in NE expression supporting activity in the fronto-parietal network, that is implicated in cognitive control (Robison et al., 2022; Unsworth & Robison, 2017). Similarly, others identified a link between pupil size and theta activity in cognitive control tasks (Dippel et al., 2017; Lin et al., 2018). We employed a concurrent registration of EEG and pupillometry to explore the relationship between activity within the LC-NE system and oscillatory brain activity in the WM task.

## Methods

The methods overlap considerably with our previously published data descriptor (Pavlov et al., 2022). Nevertheless, essential details are not omitted and fully described.

### Participants

Eighty-six participants (74 females, age: 20.5 ± 3.9 years) completed the task. The participants had normal or corrected-to-normal vision and reported no history of neurological or mental diseases. All the participants were Russian native speakers. None of the participants had had any disease of the nervous system, psychiatric, or hearing disorders in the past, or had reported use of any medication. Informed consent was obtained from each participant. The experimental protocol was approved by the Ural Federal University ethics committee.

We used the Annett Handedness Scale (Annett, 1970) to determine the handedness. Three participants were ambidextrous, 6 left-, and 77 right-handed. Ocular dominance was determined with three different tests: Rosenbach’s test (Rosenbach, 1903), aiming (the participants were asked to make the finger gun gesture with both hands and then aim at a self-selected target while closing one of the eyes (the other one is the dominant eye)), and hole-in-the-card (Fink, 1938). There were twenty-five participants with the left and 61 with the right dominant eye.

### Task and procedure

Before the WM task, we recorded 4 minutes of resting state EEG. The participants were seated in a comfortable chair and were asked to close their eyes and sit still. After the resting state recording, the participants were given instructions and proceeded with the WM task (see Figure 2).

**Figure 2.**
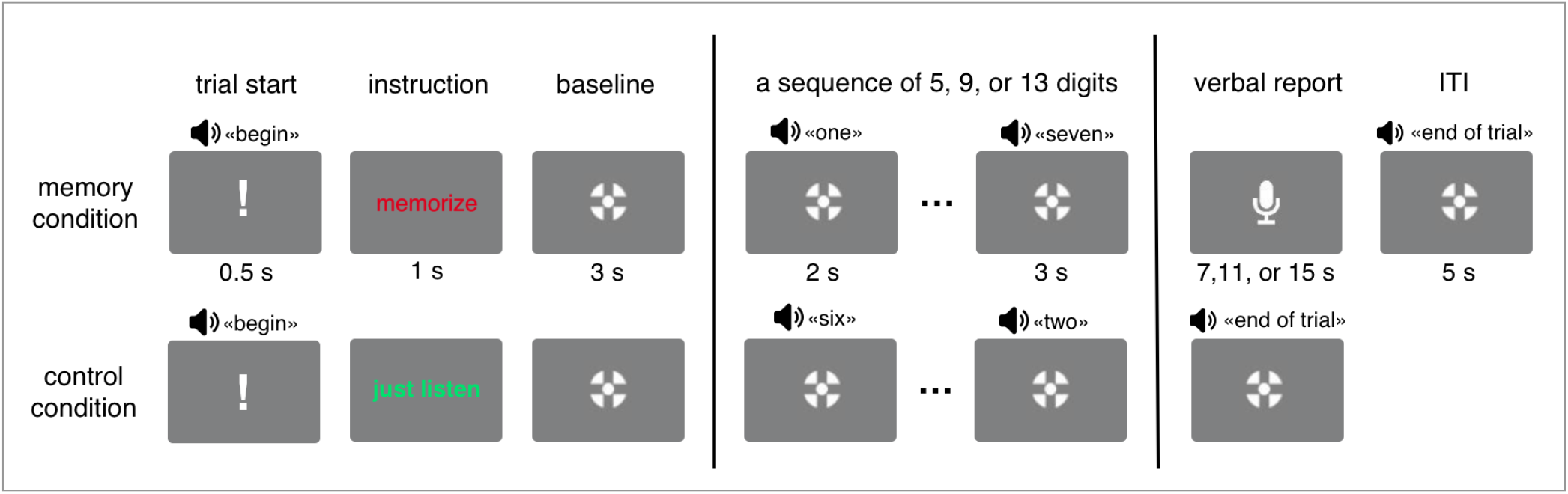
A visual representation of the experimental paradigm (not in scale). The participants were instructed to either memorize or just listen to sequences of 5, 9, or 13 digits, and then in the memory condition, they recalled the whole sequence in the serial order.

Each trial began with an exclamation mark for 0.5 s along with a recorded voice command “begin” – indicating the start of the trial. The exclamation mark was followed by an instruction to either memorize the subsequent digits in the correct order (memory condition) or to just listen to the digits without attempting to memorize them (control condition). The instruction was followed by a three-second baseline period. Then either 5, 9, or 13 digits were presented auditorily with an SOA of 2 seconds. Each of the digits from 0 to 9 was used, and the mean duration of each digit was 664 ms (min: 462 ms, max: 813 ms). The last digit in the sequence was followed by a 3-second retention interval. During the baseline, encoding, and retention intervals, participants were fixating a cross (Thaler et al., 2013) (0.8 cm in diameter) on the screen (23.6 inches in diagonal). In the memory condition, the participants were asked to recall each digit out loud in the correct order starting from the first one (i.e., serial recall). The retrieval was recorded by a computer microphone controlled by PsychoPy (Peirce et al., 2019). The participants had 7, 11, and 15 seconds for 5-, 9-, and 13-digit sequences, respectively, to recall the digits. The retrieval was followed by an intertrial interval (ITI) of 5 s. In the control condition (passive listening), the presentation of the digits and the delay period was immediately followed by an ITI of the same duration.

There were 9 blocks in total with 54 passive listening and 108 memory trials overall. Each block consisted of 3 control (one of each sequence length) followed by 12 memory (4 trials on each sequence length) followed again by 3 control trials. Memory trials within each block were presented in random order. Trial composition in a block followed the procedure from the original study by Peavler (1974). Before the main WM task, each participant completed 6 practice trials (3 passive listening and 3 memory trials). After each block, the participant had a self-paced break. After every 3 blocks, the participants took a longer break when they could take a snack and filled out a NASA-TLX questionnaire (Hart & Staveland, 1988) to self-assess the level of mental load, physical load, temporal demand they experienced, their perceived overall performance, effort, and frustration during the preceding three blocks of the task. The average duration of the experiment was 2 hours and 10 minutes.

After the whole experiment, the participants went through a structured interview about their memory strategies. The experimenter first asked an unrestricted question about whether the participant used a certain strategy. Then they were asked several more specific questions about chunking, rehearsal, memorizing only a limited number of digits at the end or the beginning of the sequences, and whether they used any visualization technique. We were able to extract 85 verbal structured reports (1 missing) on used memory strategies.

The overall accuracy in the recall was calculated as a partial score: the number of digits correctly recalled in the correct position independently of the sequence length. Thus, if the verbal response for the sequence of 416058972 was 40615897, the partial score would be 6.

The distance to the monitor was 80 cm. The loudspeakers were placed on the sides of the monitor. The measured loudness of the digit sounds was 70 dB SPL. The luminance in the room was set at 380 lux.

### Electroencephalography

A 64-channel EEG system with active electrodes (ActiCHamp, Brain Products) was used for the recording. The electrodes were placed according to the extended 10-20 system with FCz channel as the online reference and Fpz as the ground electrode. The level of impedance was maintained below 25 kOm. The sampling rate was 1000 Hz.

EEGLAB (Delorme & Makeig, 2004) was used for data preprocessing. Each recording was filtered by applying 1 Hz high-pass and 45 Hz low-pass filters. After this, data were re-referenced to averaged reference and the Independent Component Analysis was performed using the AMICA algorithm (Palmer et al., 2012). Components clearly related to eye movements were removed. Additionally, components that were mapped onto one electrode and could be clearly distinguished from EEG signals were subtracted from the data. The data were epoched in [-1500 3500 ms] intervals in relation to the presentation of the digits to avoid edge artifacts. Epochs still containing artifacts were visually identified and discarded.

All epochs were then converted into current source density (CSD) by means of CSD toolbox (Kayser, 2009). We used spherical spline surface Laplacian (Perrin, Pernier, Bertrand, & Echallier, 1989) with the following parameters: 50 iterations; m = 4; smoothing constant λ = 10−5 (for a detailed description of the procedure see (Tenke & Kayser, 2005))).

Time-frequency analysis was performed on the preprocessed single trial data between 1 and 45 Hz with 1 Hz steps using Morlet wavelets with the number of cycles varying from 3 to 12 in 45 logarithmically spaced steps for each participant and condition, separately. The analysis time window was shifted in steps of 20 ms. Spectral power was baseline-normalized by computing the percent change of the power with respect to the [-1000 to 0] ms time interval, which corresponded to the last second of the baseline fixation before the presentation of the first digit in a sequence. The time-frequency analysis was performed by means of the Fieldtrip toolbox (Oostenveld et al., 2011).

For EEG, two frequency-bands and regions of interest (ROI) were identified: frontal midline theta (4-8 Hz, channel Fz) and posterior alpha (9-14 Hz, channels P1, P2, Pz, P3, P4). Fz was selected because it has been used for quantifying frontal midline theta in most previous studies (Pavlov & Kotchoubey, 2020). We included more channels for quantification of alpha because of a more widespread topography as revealed by cluster-based permutation tests comparing control and memory conditions (see Figure 5).

Three EEG recordings were excluded because of technical failure and 19 recordings were excluded due to an experimenter error resulting in the misplaced electrode cap being incompatible with 10-20 system electrode layout. Thus, we are reporting the analysis based on 65 EEG recordings.

### Pupillometry

Pupillometry was recorded with Pupil Labs wearable eye-tracker (Pupil Lab GmBH, Germany) with 120 Hz sampling rate. One-point calibration preceded each recording. For preprocessing, first, we selected data from the dominant eye. Next, the data points identified by Pupil Player algorithm as having less than 80% confidence score were replaced with missing values (coded as ‘NA’) for further interpolation. Then, the data were broken into epochs corresponding to the time intervals starting 2 seconds before presentation of the first digit in the sequence and ending 3 seconds after the presentation of the last digit in the sequence.

Further processing was conducted using Gazer package for R (Geller et al., 2020). First, the median absolute deviation was used to identify rapid pupil size changes (see Geller et al. 2020 for details). Values above 16 median absolute deviations were replaced with missing values. Then, the blinks (detected through the Pupil Labs algorithm) were appended with time intervals -100 before and +100 ms after each blink and replaced with missing values. Trials containing more than 50% of the missing values were removed from the analysis. Participants with less than 6 remaining trials in any condition were excluded from the analysis. In the end, the missing values in the remaining trials were linearly interpolated and smoothed with 5 points moving average.

The resulting pupil size was baseline normalized by the subtraction of the mean absolute value in the interval of 2 seconds before the presentation of the first digit in the sequence. Then the data were downsampled to 10 Hz.

Two pupillometry datasets were excluded because of technical failure with the recording software, and the artifact correction and rejection procedure resulted in the exclusion of additional 11 participants. Thus, 73 participants were included in the analyses.

### Statistics

For exploratory analysis of EEG, non-parametric cluster-based permutation tests as implemented in Fieldtrip toolbox (Oostenveld et al., 2011) for MATLAB. We applied the tests with 5000 permutations in spectral power averaged in 2-second intervals after the presentation of the digits with at least 2 channels and alpha of 0.05 threshold for forming a cluster.

The remaining statistical analyses were conducted in R (v. 4.2.0). For the first repeated-measures ANOVA of EEG and pupillometry data, relative spectral power and pupil size were averaged in 2-second intervals corresponding to encoding and maintenance of individual digits. This analysis included factors Load (13 levels) and Task (2 levels: control vs memory). For the second ANOVA the data were averaged in the last 3 seconds before recall corresponding to the retention interval after the last digit in the sequence. This analysis included factors Task and Sequence (3 levels: 5, 9, and 13 digits). Greenhouse-Geisser adjusted degrees of freedom are reported were applicable.

When appropriate, we tested the lack of difference between conditions by calculating Bayes Factors (BF) using a Bayesian t-test with default priors, where BF_01_ – evidence in favor of null hypothesis and BF_10_ – evidence for the alternative hypothesis. BF were always reported to be above 1 (with varying the indices). For this analysis, we used BayesFactor package for R. In the regression analysis, all variables were first centered and then scaled by dividing by the standard deviations to produce standardized beta coefficients.

## Results

### Behavior

#### Accuracy

The number of correctly recalled digits significantly decreased as the length of the sequence increased (main effect of Sequence (5- vs 9- vs 13-digit sequences): F(1.69,143.40) = 60.32, p < .001, 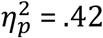). Mean (±SD) number of recalled digits was 4.62 ± 0.31 for 5-digit, 4.18 ± 1.18 for 9-digit and 3.57 ± 1.13 for 13-digit sequences. Recall accuracy decreased with load in every sequence (main effect of Load for 5-digit (F(2.42,205.34) = 59.36, p<.001, 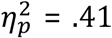), 9-digit (F(3.41,289.52) = 606.36, p < .001, 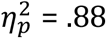), and 13-digit (F(2.46, 208.82) = 549.86, p < .001, 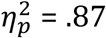) sequences). Mean accuracies for each level of load and sequence length are depicted in Figure 3A. A pairwise comparison between levels of load in each sequence length showed highly significant differences (p < 0.001) for almost all pairs. For the 5-digit sequence, there were no differences in recalling between the 3rd and 4th, 3rd, and 5th digits (p = 0.157). The difference between the 12th and 13th digits was not significant (p = 0.153) in the 13-digit sequences. These results indicate preserved recency effect (significantly better recall of the last digit in the sequence than of the second to last) for 5- and 9-digit sequences but not for 13-digit sequences.

**Figure 3.**
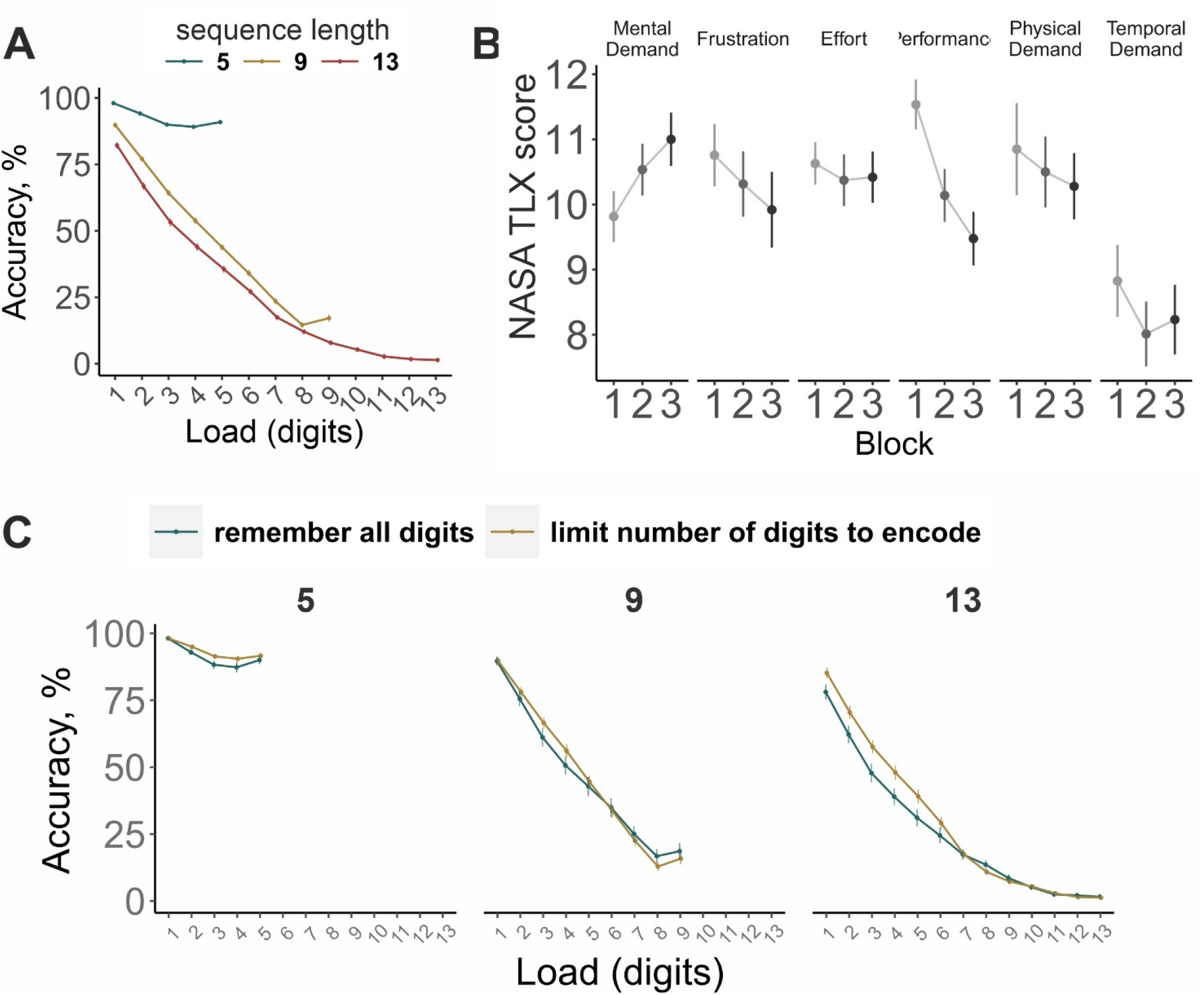
(A) mean accuracy in recalling digits separated by sequence length (5, 9, 13 digits), (B) mean scores on 6 NASA-TLX subscales on 20 point scale (how (1) mentally demanding, (2) frustrating, (3) effortful, (4) difficult to perform, (5) physically demanding and (6) temporally demanding the task was), which participants gave after each of the 3 large blocks of trials. (C) Comparison of two memory strategies: to remember all presented digits in the sequence or intentionally remember only a limited number of digits and ignore the rest. Error bars depict Standard Error of the Mean (SEM).

#### Subjective task difficulty

Time on the task (a potential source of fatigue) did not influence overall perceived cognitive load (main effect of Block: F(2, 170) = 1.80, p = .169, 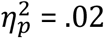, Block x Scale interaction: F(6.92, 587.80) = 1.76, p = .094, 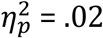; see Figure 3B).

#### Memory strategies

A descriptive analysis of the strategies revealed that 74/85 participants used chunking, 63/85 employed some visualization strategy (e.g., mapping the digits on a dial pad, table, visualizing the sequence, color them), 75/85 rehearsed at least some of the digits, while 66/85 rehearsed the whole sequence adding each newly presented digit to the cumulative rehearsal, 49/85 used a strategy to memorize only restricted number of the first digits in the sequence, and 31/85 reported attempting to memorize a limited number of the last digits in the sequence.

The highly disproportionate group sizes did not allow us to explore the effectiveness of most strategies with enough statistical power. However, we could compare the two strategies: (1) to intentionally remember only a limited number of the first digits in the sequence and ignore the rest and (2) to continue trying to encode and maintain every presented digit in the sequence. The difference in recall performance was not significant between the groups using these two strategies (main effect of Group: F(1, 83) = 1.13, p = .29, 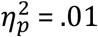; see Table S1). But the analysis of the accuracy at each level of load in three sequence lengths separately revealed a significant interaction between Load and Group in the 13-digit sequences (see Figure 3D and Table 1) and 9-digit sequences (p = 0.051, conventionally non-significant) but not in the 5-digit sequences. These results indicate better recall for the digits before the 7^th^ in the 13-digit sequences in the group of participants who reported stopping encoding digits before the end of the sequence than in the other group.

**Table 1.**
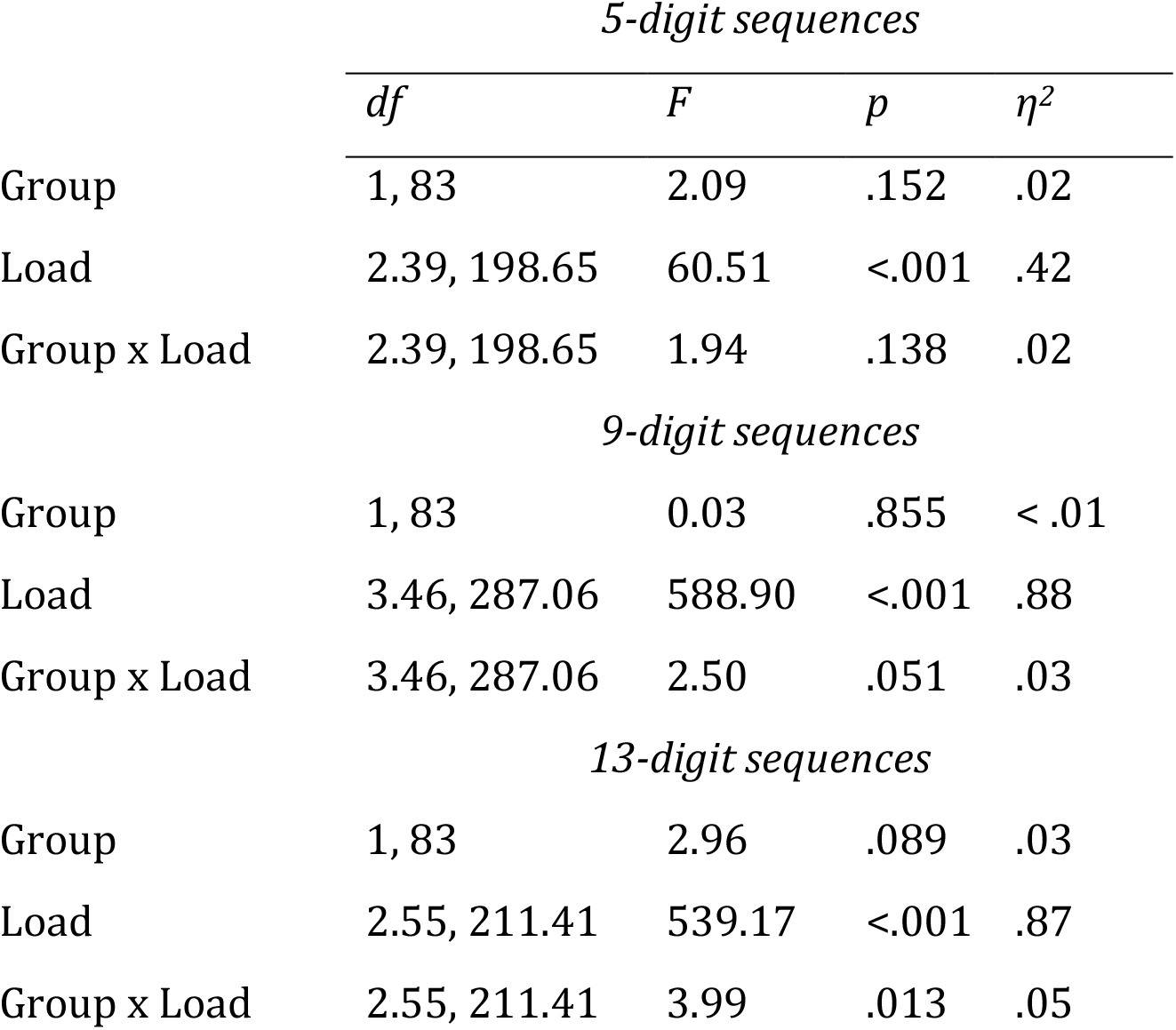
The effect of strategies on recall accuracy

| *5-digit sequences* |  |  |  |  |
| --- | --- | --- | --- | --- |
| | *df* | *F* | *p* | $\eta^2$ |
| Group | 1, 83 | 2.09 | .152 | .02 |
| Load | 2.39, 198.65 | 60.51 | <.001 | .42 |
| Group x Load | 2.39, 198.65 | 1.94 | .138 | .02 |
| *9-digit sequences* |  |  |  |  |
| Group | 1, 83 | 0.03 | .855 | < .01 |
| Load | 3.46, 287.06 | 588.90 | <.001 | .88 |
| Group x Load | 3.46, 287.06 | 2.50 | .051 | .03 |
| *13-digit sequences* |  |  |  |  |
| Group | 1, 83 | 2.96 | .089 | .03 |
| Load | 2.55, 211.41 | 539.17 | <.001 | .87 |
| Group x Load | 2.55, 211.41 | 3.99 | .013 | .05 |

A comparison of the strategy of giving priority to the last few digits in the sequence and the strategy of remembering all digits revealed that the former strategy did not yield any benefits.

The strategy might have developed throughout the experiment. To compare our results more directly with the original study and rule out the possible effect of more effective memory strategies developed throughout the session, we included only one block of trials in the analysis (akin to the original study by Peavler). The effect of improved recall for the first digits in the 13-digit sequence was also observed in the first block of trials (Group x Load: *F*(6.29, 521.98) = 2.85, *p* = .009, 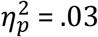). This strategy was successful for the recall of the first 5 digits in 13-digit sequences (p-value for the 5^th^ digit 0.0535 and the rest before the 5th p < 0.05).

### Pupillometry

#### Memory condition

Our study was designed to resolve the contradiction between the initial work by Peavler and the follow-up replication by Granholm et al. concerning the temporal trajectory of pupil size changes with increasing WM load.

The pupil dilated in response to the presentation of the first 6 digits, and, after plateauing from 7- to 10-digit load, declined reaching the baseline level after the presentation of the last 13^th^ digit (Figure 4). This observation is supported by a significant Load (F(1.94, 139.89) = 10.97, p < .001, 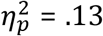) and Task effects (F(1, 72) = 126.26, p < .001, 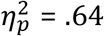), and their interaction (F(2.30, 165.91) = 59.19, p < .001, 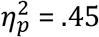).

**Figure 4.**
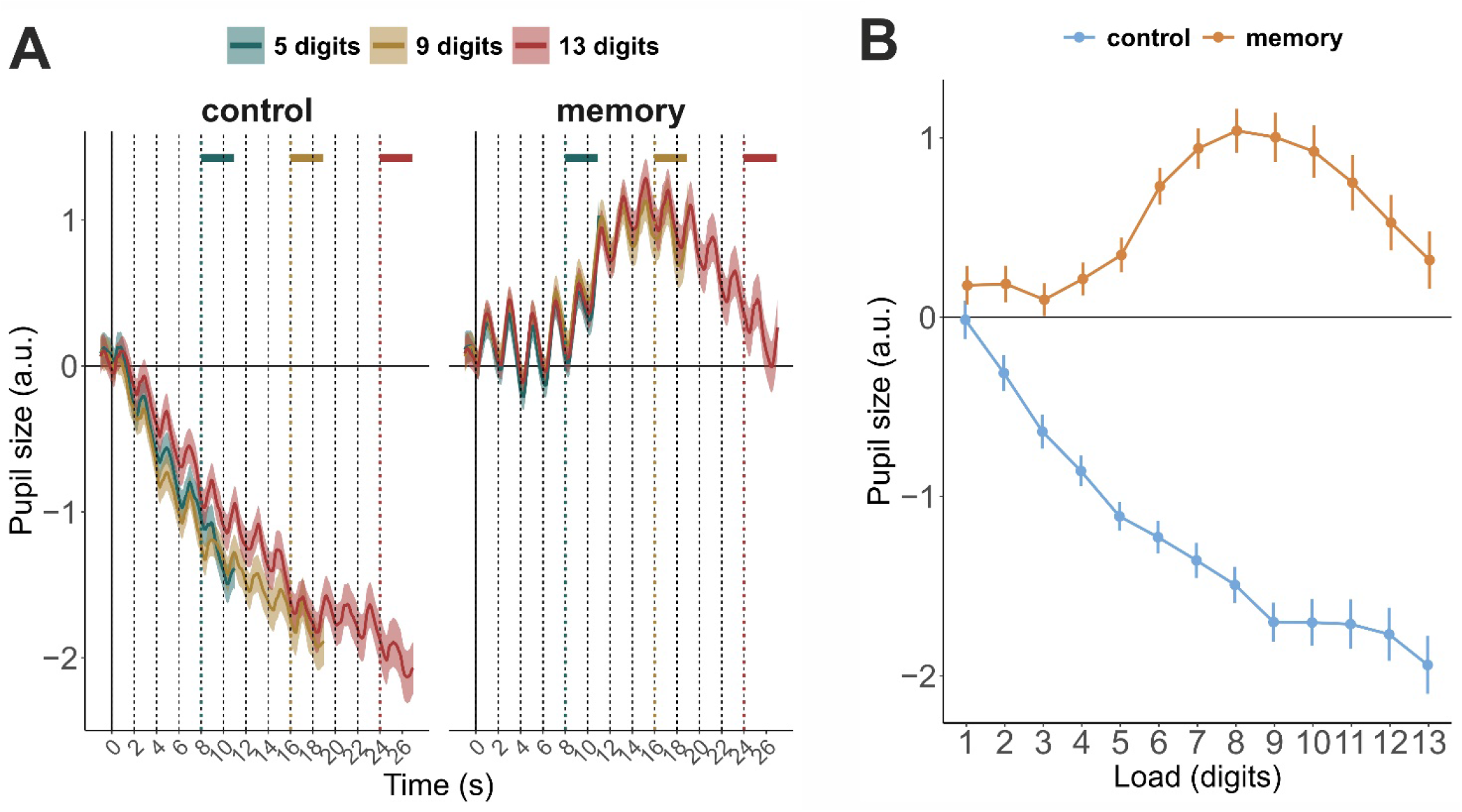
Pupillometry results. (A) pupil size changes in time in 2 conditions and 3 sequence lengths. Dashed vertical lines mark the presentation of single digits in the sequences. Colored horizontal bars mark the retention interval after presentation of the last digit in the sequences. (B) pupil size averaged in the two second intervals after presentation of the digits.

On average, after the presentation of the last digit in the 13-digit sequences, pupil size was no longer different from baseline level. This effect is supported by one-sample t-tests of average pupil size during the retention interval (i.e., the last 3 seconds before recall) (5-digit sequence: t(72) = 2.07, p = 0.042, BF_01_ = 1.04; 9-digit: t(72) = 4.33, p < 0.001, BF_10_ = 411; 13-digit: t(72) = 1.09, p = 0.281, BF_01_ = 4.42). Additionally, we repeated the analysis employed by Granholm et al. that showed a non-significant pupil dilation in the last second before the recall. This analysis yielded even stronger evidence for the null effect after presentation of the last digit in the 13-digit sequences but not after presentation of the last digits in the 5 and 9-digit sequences (5-digit sequence: t(72) = 3.04, p = 0.003, BF_10_ = 8.64; 9-digit: t(72) = 3.79, p < 0.001, BF_10_ = 74.4; 13-digit: t(72) = 0.36, p = 0.718, BF_01_ = 7.29).

When we left only one block of trials to make the number of trials comparable to the original study, the dynamics was similar but not identical (see supplementary Figure S1). Although pupil started steeply decreasing after the peak, it remained above the baseline level till the end in the 13-digit sequence condition (after exclusion of 6 participants with less than 2 trials in the condition, one-sample t-test for the last second before recall: t(66) = 2.178, p = 0.033, BF_10_ = 1.2). In all other blocks of trials, if analyzed separately, the test did not yield significance and the BF_01_ were above 2.

The strategy type did not significantly affect pupil size in any of the sequence lengths (see Figure S2).

#### Control condition

As can be seen in Figure 4, in the control condition (passive listening), pupil size declined after the presentation of the first digit and continued the trend till the end of the sequence. The difference between the control and memory conditions was significant for all levels of load.

### Electroencephalography

#### Spectral power in theta and alpha frequency bands

To explore neural correlates of WM loading and overloading, we analyzed the EEG. As can be seen in Figure 5, after averaging over sequence lengths, theta and alpha activity strongly differed between the control and memory tasks (main effects of Task, Load, and the Task x Load interaction; see Table 2). As can be seen in Figure 5A, presentation of the digits during the passive listening task elicited short-lived theta bursts but a continuous increase in theta power was only observed in the memory condition. Theta activity was not significantly different between the control and memory conditions when only one or two items have been presented (Figure 5C). Similarly, we found no effect of the task on alpha activity during the first four digits presentation. These observations are also supported by data-driven cluster-based permutation tests that revealed no significant clusters for the first two digits in the theta and alpha frequency bands (Figure 5B).

**Figure 5.**
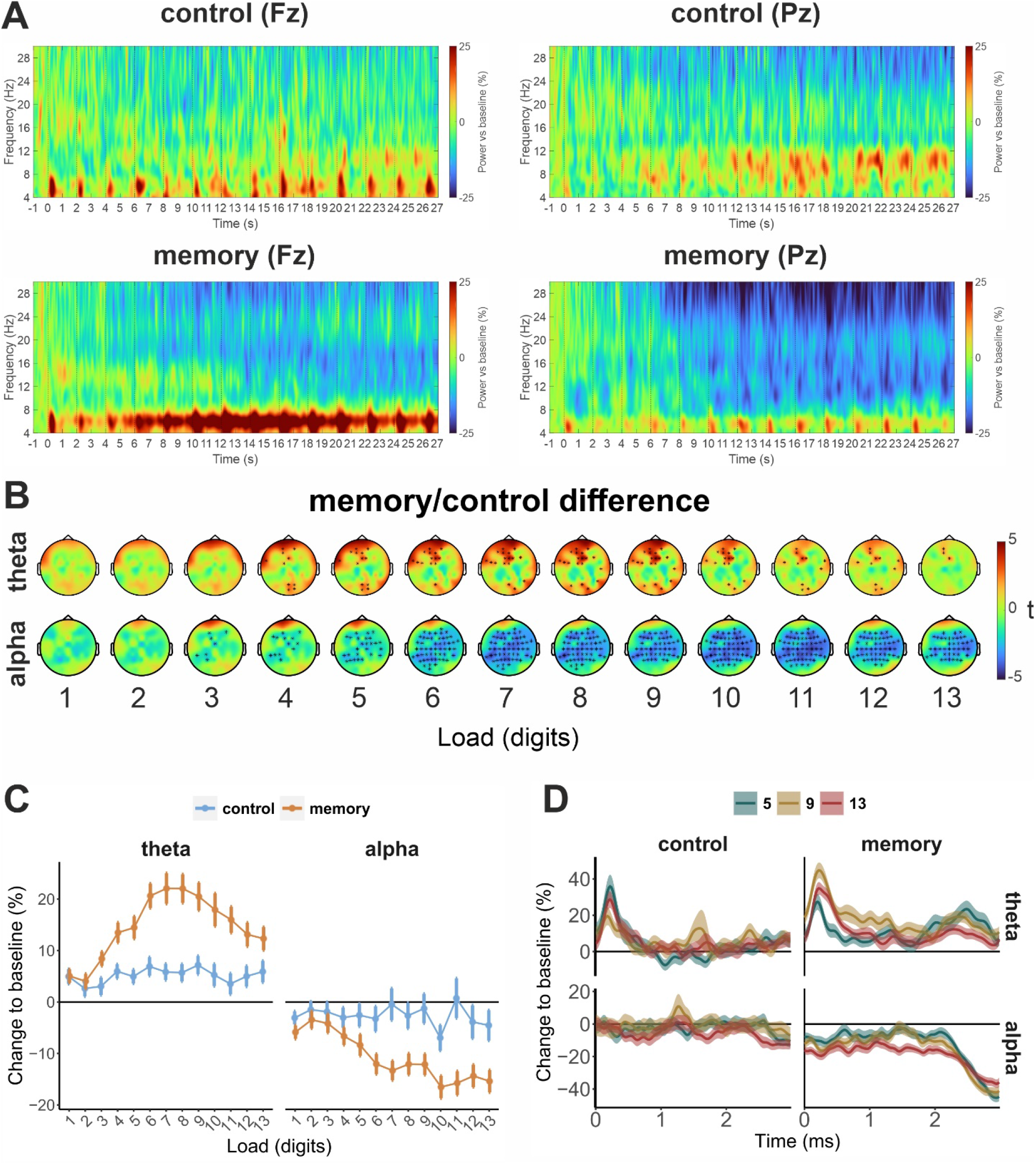
(A) Time-frequency maps at representative channels (Fz and Pz). Alpha and theta power over time in two tasks in 13-digit sequences over time. (B) Topography of the difference in control and memory tasks. Stars mark electrodes with significant effect of Task. (C) Load effects. The spectral power of alpha (9-14 Hz) and theta (4-8 Hz) activity is averaged in two-second time intervals corresponding to encoding and maintenance on individual digits averaged over all sequence lengths. (D) The effect of sequence length on spectral power in the retention interval (three seconds before recall).

**Table 2.** Load and task effects on oscillatory brain activity

|  | theta |  |  |  | alpha |  |  |  |
| --- | --- | --- | --- | --- | --- | --- | --- | --- |
| | *df* | *F* | *p* | $\eta^2$ | *df* | *F* | *p* | $\eta^2$ |
| **Load** | 5.72, 365.89 | 11.82 | <.001 | 0.156 | 4.62, 295.49 | 5.41 | <.001 | 0.078 |
| **Task** | 1, 64 | 17.91 | <.001 | 0.219 | 1, 64 | 11.21 | 0.001 | 0.149 |
| **Load x Task** | 6.04, 386.49 | 7.05 | <.001 | 0.099 | 4.35, 278.63 | 4.40 | 0.001 | 0.064 |

Theta activity during the memory task was changing with increasing memory load. There was a general upwards trend up to 7-digit load. The trend reversed after the peak till the presentation of the last 13^th^ digit in the longest sequence when theta power approached the level observed at 4 digits load (t(64) = 0.531, p = 0.597, BF_01_ = 6.42). The analysis of consecutive changes in alpha activity with load showed that alpha significantly decreased from 3 to 4 digits load, continued decreasing up to 6-digit load, and then another drop in power occurred after the 9^th^ digit (see Tables S2-S7 for pairwise comparison results).

Theta and alpha activity after the presentation of the last digit (3 s) – during the retention interval – did not depend on the sequence length, but in the memory task, alpha suppression and theta enhancement were stronger than in the control task (the main effect of Task; see Table 3 and Figure 5D). Thus, the average amplitude of alpha and theta activity after the presentation of the last digit in 5, 9, and 13-digits sequences was comparable. The memory strategy to limit the number of digits did not modulate the temporal dynamics (no Group main effect or interactions with Task and Load).

**Table 3.**
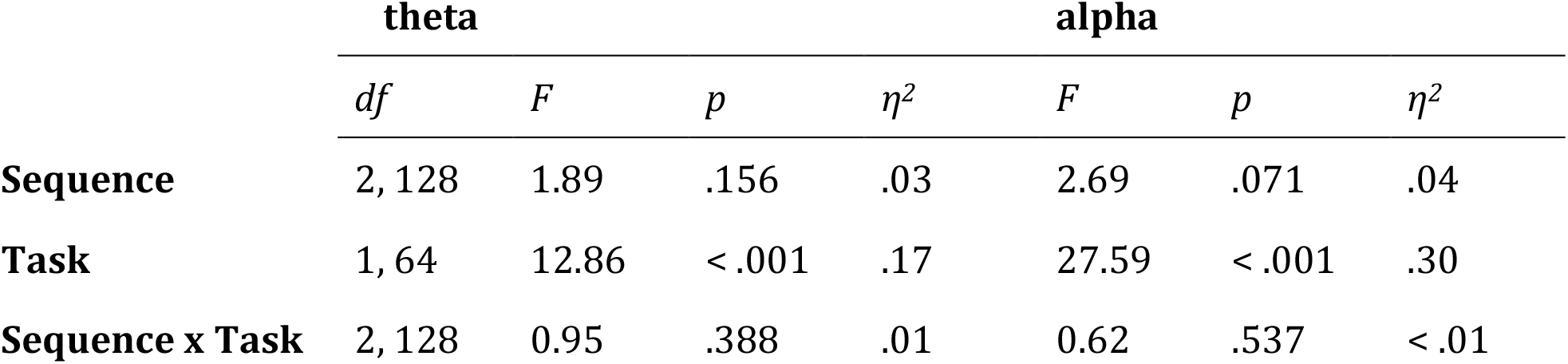
Averaged in the retention interval (3 seconds after presentation of the last digit in the sequence)

**Table 3.** Regression table for 5-digit sequence recall accuracy

| Predictors | $\beta$ | CI | t | p |
| --- | --- | --- | --- | --- |
| Pupil size | 0.06 | -0.22 – 0.34 | 0.41 | 0.681 |
| Alpha power | -0.17 | -0.44 – 0.11 | -1.23 | 0.224 |
| Theta power | 0.29 | 0.01 – 0.56 | 2.08 | **0.042** |

**Table 4.** Regression table for 9-digit sequence recall accuracy

| *Predictors* | $\beta$ | *CI* | *t* | *p* |
| --- | --- | --- | --- | --- |
| Pupil size | 0.37 | 0.12 – 0.63 | 2.96 | **0.005** |
| Alpha power | -0.23 | -0.47 – 0.00 | -1.98 | 0.053 |
| Theta power | 0.02 | -0.24 – 0.28 | 0.12 | 0.905 |

**Table 5.** Regression table for 13-digit sequence recall accuracy

| *Predictors* | $\beta$ | *CI* | *t* | *p* |
| --- | --- | --- | --- | --- |
| Pupil size | 0.25 | -0.04 – 0.53 | 1.70 | 0.095 |
| Alpha power | -0.25 | -0.53 – 0.03 | -1.78 | 0.081 |
| Theta power | 0.07 | -0.21 – 0.35 | 0.52 | 0.605 |

#### Individual differences in physiological functioning and recall accuracy

First, we investigated the relationship between average values over all sequences and digits (except the retention interval) of spectral power and pupil size and behavioral performance. The reliability estimates (Cronbach’s alpha) and other descriptive statistics of the studied variables are reported in Table S8.

Pupil dilation had a positive association with behavioral performance (Figure 6A). For a visual depiction of the temporal dynamics of pupil dilation in participants with different memory capacity, we divided the sample into high and low performing groups based on median split. The previously described triphasic pattern of the changes in pupil size was not dependent on behavioral performance and remained similar in high and low performing individuals (see Figures 6C). Figure 6C suggests that participants with better accuracy scores had stronger pupil dilation during memorizing digits around the plateau (7-10^th^ digits) but the Group x Load interaction was not significant in the memory task (*F*(1.86, 132.02) = 1.85, *p* = .164, 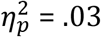). But the high performance group showed stronger pupil constriction in the passive listening condition (Group x Load (*F*(2.48, 176.21) = 3.10, *p* = .037, 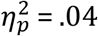) in the control task and Group x Load x Task (*F*(2.36, 167.69) = 5.73, *p* = .002, 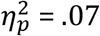)) .

**Figure 6.**
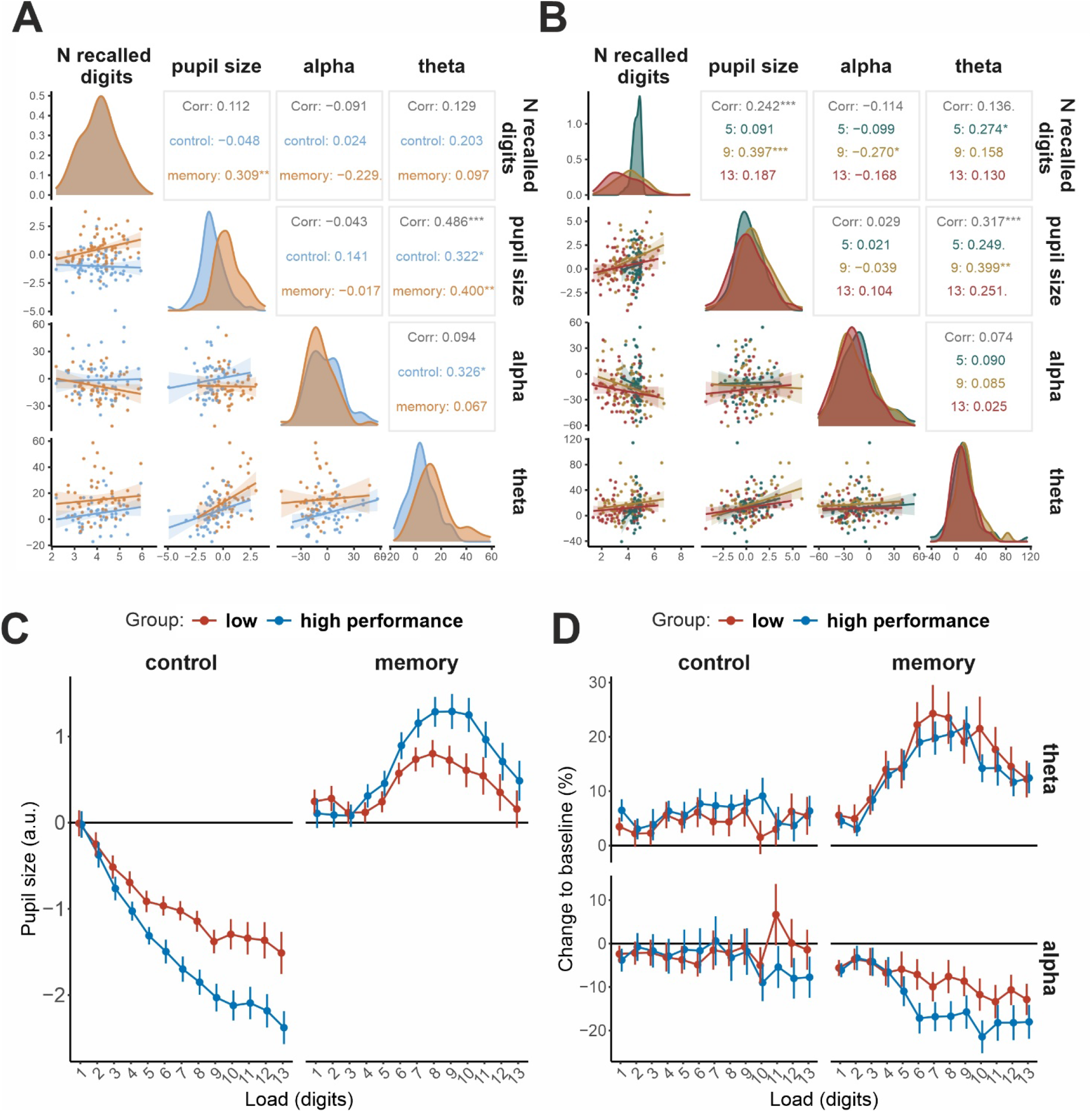
Correlation matrix with scatter plots. (A) Correlations of physiological responses averaged over all time points except retention interval; (B) Same, but for the retention interval. A comparison between high and low performance groups in their (C) pupil size and (D) spectral power.

Then we estimated individual peaks of pupil dilation using data from the 13-digit sequences. For this analysis, pupil size was first averaged in two-second intervals corresponding to encoding and maintenance of each digit in the sequence in every trial separately, peaks were identified on a single trial level, and then individual digit-at-peak dilation was calculated as an average over trials. Digit-at-peak dilation was 7.28 ± 1.62 digits on average. The peak dilation correlated with behavioral accuracy in 5-, 9-, and 13-digit sequences (r(71) = 0.42, 0.40, and 0.27, p < 0.001, <0.001, and 0.019, respectively) and overall accuracy across all conditions (r(71) = 0.37, p = 0.001).

The average power of alpha activity correlated negatively with the average number of recalled digits (independently of the length of the sequence), while no correlation was found between theta power and recall accuracy. Theta power and pupil size demonstrated a positive correlation during memory (r(63) = 0.400) as well as in the control task (r(63) = 0.322) (Figure 6A). To rule out the potential effect of event-related potentials (broadband changes in spectral power due to non-genuine oscillations) on the pupil-theta correlation, we limited the analysis window for theta activity to the second second after the digit onset. This manipulation did not affect the character of the relationship (memory: r(63) = 0.404; control task: 0.382; see Figures S3 and S4).

To assess the relationship between physiological indices of cumulative WM load and behavioral accuracy we included in the analysis the retention interval (3 seconds after presentation of the last item in the sequence) of each sequence length separately. In the 9-digit condition, we found a significant relationship between both pupil size (r(71) = 0.397) and alpha activity (r(63) = - 0.270) correlated with the number of recalled digits. In the 5-digit condition, theta activity during the retention interval was positively associated with recall accuracy (r(63) = 0.274). To take a closer look at this relationship, we explored correlations between accuracy in recall of the digits in the first 5 serial positions and corresponding theta power (e.g., accuracy in memorizing 1^st^ digit in 13-digit sequences and corresponding theta power in the same condition). This analysis revealed significant correlations between theta power during encoding and retention of the 1^st^ (r(63) = 0.27, p = 0.0281), and 5^th^ (r(63) = 0.28, p = 0.0226) digit in the 5-digit sequence but not for the 2^nd^ (r(63) = 0.11, p = 0.3835), 3^rd^ (r(63) = 0.18, p = 0.1412), and 4^th^ (r(63) = 0.16, p = 0.1947) ones. The same analysis for 9-digit and 13-digit sequences did not yield any significant correlations. This analysis suggests that this correlation is unique to 5-digit sequences, not to the first five digits in any sequence.

To estimate a joint effect of the oscillatory brain activity and pupil dilation (during the retention interval) on recall accuracy, we also applied multiple regression for each sequence length separately. The results of this analysis only confirmed the correlation analysis results showing that the predictors have largely independent contributions to the recall accurac The perceived mental effort might have affected the hypothesized indices of mental effort: pupil size and theta power. However, none of the NASA-TLX scales showed a significant relationship with the physiological variables (see Figure S5). Participants who perceived the task as less demanding or applied more effort did not recall a larger number of digits.

## Discussion

### Pupil size temporal dynamics

To resolve the debate between previous research on the pupil dilation dynamics under working memory overload (Granholm et al., 1996; Peavler, 1974), we adopted the digit span task employed previously by Peavler (1974) and later by Granholm et al. (1996) where participants either memorized or passively listened to sequences of 5, 9, or 13 digits. Increasing sample size and number of trials was crucial to making a convincing argument and to achieving greater precision and statistical power. In result, in the passive listening control condition, the pupil started constricting immediately after the presentation of the first digit in the sequence as compared to the memory condition where pupil size dynamics can be described by the triphasic model (Figure 1): linear increase from serial positions 1 to 6, a plateau from 7 to 10, and then a decline from 11 to 13, with pupil size no longer different from baseline after the presentation of the 13^th^ digit. The downward trend in pupil size appears to develop after reaching the predicted individual digit span of 7 digits. This temporal pattern was neither related to behavioral performance nor memory strategy. Thus, under cognitive overload – when the number of memory items exceeds WM capacity – the pupil starts constricting, but only after a plateau. The observed data replicates the pattern of the results documented by Granholm et al. (1996).

Pupil dilation during performance of working memory tasks has been previously associated with the exertion of effort or, in other words, allocation of mental resources to the task at hand (Beatty & Kahneman, 1966; Kahneman & Beatty, 1966). Applied effort and working memory load are expected to increase with adding new items to the sequence. However, the initial uptrend was replaced by a plateau – small insignificant changes in pupil size – between 7 to 10 digits. The plateau phase can be explained by the participants maintaining a limited number of digits in the focus of attention for several more seconds after reaching their individual capacity. Then, when they were no longer able or refused to allocate the resources to maintain attention, the pupil started constricting. Our results support previous findings that the initial increase in pupil size is not indefinite and overtaxing individual cognitive resources leads to pupil stopping dilating (Unsworth & Miller, 2021).

One possible source of variability in the temporal dynamics of pupil dilation and constriction is individual differences in behavioral performance. The absolute peak of pupil dilation varied individually depending on behavioral performance. In absolute values, the dilation continued up to about 8-digit load on the group level. Measured individually, we observed the average peak at 7.27 digits. Both of these numbers are larger than the expected digit span (i.e., 7 – the average digit span of an individual (Miller, 1956) or 6.74, as taken from normative data for young adults aged 20-29 (Woods et al., 2011)). But these results are in line with a previous study where the peak was reached at 7.7 digits (Johnson et al., 2014). Similarly, the peak was observed at 7-item load when words were used as stimuli (Unsworth & Miller, 2021). It is also in line with a visual working memory study, where the pupil size peak was observed at 5 items load (colored squares) and similarly exceeded the maximum number of items that an individual could hold in WM (i.e., Cowan’s k of 4) (Unsworth & Robison, 2015, 2018). We did not measure the digit span with the traditional procedure (Wechsler, 1955) but recall accuracy in the used version of the task must be a correlate of the traditional measure. In a previous study that measured digit span directly, the span was 7.2 and it correlated with the digit-at-peak pupil size (r = 0.38) (Johnson et al., 2014). We showed that, although digit-at-peak pupil size correlated with behavioral accuracy on average, low- and high-capacity individuals showed similar temporal dynamics. On the contrary, in the experiment of Tsukahara et al. (2016; the longest sequence of 12 letters), pupil constricted after peaking at the 8^th^ item in the low-performance group, but pupil plateaued after the 9^th^ item in the high-performance group. The exact reason for the discrepancy between the two sets of results remains unclear; potential factors of the disagreement are differences in motivation between our samples and the modality of the stimuli (visual in Tsukahara et al.). Nevertheless, it appears that the peak does not represent individual WM capacity, but rather an ability to maintain attention after approximately the 5^th^ digit in the sequence. Overall, our results speak for the resource allocation hypothesis account of the observed differences – the high-capacity group had a larger resource pool than the low-capacity group who could not exert more effort to memorize the later items in the sequence.

Adopted memory strategies might affect the temporal dynamics and peak pupil size. Many participants reported limiting the number of digits to retain as their strategy. We can hypothesize that the participants on average could keep about 7 digits in memory. Perhaps they continued maintaining the first 7 digits while the new digits were presented in the 9-digit and 13-digit sequences and tried to suppress the new stream of information. The new digits not fitting to memory were playing the role of distractors resulting in diminished performance in memorizing longer sequences. The strategy to maintain only a limited number of digits led to increased accuracy in the longest 13-digit sequences for the first 5 digits. But this advantage was not reflected in pupil size, suggesting that this strategy did not come with any additional effort during both encoding the earlier digits and suppression of the later digits in the sequence.

Another question widely discussed in the literature is the relation between cognitive effort and pupil diameter. We found that the relationship between pupil size and subjectively perceived cognitive load was not significant. Although some previous studies of physical effort (Zénon et al., 2014) and mental workload (the average of all NASA-TLX scales) in a set of cognitive tasks including mental arithmetic (Recarte et al., 2008) found the correlation, pupil dilation has not always been associated with self-reports in the study of listening effort (Wendt et al., 2016; Zekveld & Kramer, 2014; Zekveld et al., 2011). The lack of significant correlation in our study can possibly be attributed to the roughness of the measure: we asked the participants to make the self-assessment after a large block of trials containing both passive and active tasks, with and without overload.

### Theta spectral power temporal dynamics

Application of the digit span paradigm with a sequential presentation of the digits along with a sufficiently long SOA allowed us to conduct exploratory analyses of oscillatory brain activity during the task. We found that frontal midline theta power was linearly tracking the WM load up to the 6-digit load which was followed by a plateau from 7 to 9 digits, and then a decrease. Unlike pupil size, theta never reached the baseline level. With a typical number of items in verbal WM studies of 5, the first phase of the temporal trajectory – the linear modulation of theta power by an increasing number of items for retention – has been reported repeatedly (Pavlov & Kotchoubey, 2022). Paralleling the observed plateau effect, our previous verbal WM study failed to observe a difference in expression of theta activity between 5, 6, and 7 letters load (Pavlov & Kotchoubey, 2021). In that study, we hypothesized that theta attained plateau at this rather high level of load. Assuming that letter span is smaller than digit span, in the current study, this hypothesis has found support. The last phase of the trajectory – the overload – has never been explored in EEG research. Here, we showed that, similarly to pupil size, increasing WM load produced first linear increase then a plateau, and then a drop in spectral power of theta activity.

Frontal theta activity has been closely associated with cognitive control (Cavanagh & Frank, 2014; Pavlov & Kotchoubey, 2021; Sauseng et al., 2010). Cognitive control may serve multiple functions in the digit span task: it can be involved in sustained attention to actively maintain information in WM (Oberauer, 2002), interfacing with long-term storage, counteracting interference between items in memory (Berman et al., 2009; Oberauer et al., 2018), shifting of attention between memory items (Camos & Barrouillet, 2014), and updating of the content of memory with newly added items. On the one hand, theta power was shown to increase with WM load supporting the role of theta activity in executive control. On the other hand, in the group of participants who chose the strategy to memorize only a selected number of digits in sequences, theta did not respond to the demand to ignore newly presented items which can be considered as distractors. This finding may suggest a more limited role of theta activity in suppressing interference from the distractors. Alternatively, suppressing distractors in our version of the task is not very effortful and does not drain executive control resources. Perhaps, maintenance and continuous rehearsal of already encoded digits are the main consumers of the resource. Additionally, the digit span task may rely on verbal storage capacity more than executive control ability.

The need for cognitive control is presumably the highest on the edge of individual capacity limits. We can hypothesize that overloading memory capacity destroys the networks supporting the control and does not allow any more refreshment of memory traces or counteracting interference. These effects were evident in behavioral performance – presentation of longer sequences did not lead to the recall of a larger number of digits. However, in our previous study (Pavlov & Kotchoubey, 2020), the relationship of theta power with behavioral performance in conditions with no excessive demand on executive control (simple retention task as compared to a task involving mental manipulations with memory content) was found to be not significant. In the current study, theta did not correlate with overall recall accuracy either.

Outside of the cognitive control framework, an influential model suggests that frontal theta coordinates the sequential reactivation of WM items each of which is represented by a different theta phase (Lisman & Jensen, 2013). Thus, theta contributes to the storage of temporal order information (Hsieh & Ranganath, 2014). The model predicts that increasing set size would lead to a more complex temporal relationship between memory items that would in turn increase theta amplitude (Hsieh & Ranganath, 2014). A critical test for this theory would be an observation that one-item WM load does not engage theta activity – the binding function of theta, in this case, is not required. The analysis using the second half of the retention interval after the presentation of the digits (e.g., after elimination of evoked activity) indeed showed a lack of theta increase as compared to baseline. Independent of the time window used for averaging, theta activity in response to the first two digits in a sequence did not differentiate between memory and control condition. Only presentation of the third item in the sequence elicited a reliable increase of theta power above baseline level. Thus, theta activity only comes into play in multi-item WM tasks with strong temporal structure (e.g., in verbal WM tasks where sequential ordering occurs naturally even with simultaneous presentation of the memory items (Gmeindl et al., 2011; Guida et al., 2020)).

### Alpha spectral power temporal dynamics

Although some studies reported alpha reaching an asymptote (either increasing or decreasing) at 5 item load (Bashivan et al., 2014; Scheeringa et al., 2009), this was not the case in our study. Unlike theta activity, alpha showed a pattern of suppression that was maintained until the end of the longest sequence presentation. The apparent difference in the temporal dynamics between theta and alpha activity suggests different functional roles of the rhythms in WM.

Alpha suppression during retention has been associated with the direction of the attention spotlight and the number of WM items in the focus of attention during the retention interval (de Vries et al., 2018; Fukuda et al., 2015; Schneider et al., 2019; Sutterer et al., 2021). Direct mapping of alpha activity on the WM items in the focus of attention is hardly supported by our data: despite memory overload, alpha continued decreasing and stabilized only at the end of the longest sequence of 13 digits. No current theory of WM predicts the ability to simultaneously maintain 13 memory items in the focus of attention but being able to recall only 3-4 of them. With that, accuracy in memorizing 9-digit sequences was related to the level of alpha suppression. Higher performers showed a trend of stronger alpha suppression in memorizing middle and late memory items after the 6th digit. Thus, the capacity of the focus of attention can vary between individuals and contribute to successful task performance but alpha is unlikely to be solely explained by the focus of attention account.

Experimental paradigms that require encoding of one or two items in WM are frequently used in the literature but rarely do researchers employ a passive perception control condition. A comparison of alpha activity in the memory and the control task revealed no difference between the tasks during encoding and maintenance of the first two items in the sequence. This finding also speaks against the focus of attention interpretation of the alpha activity. At least in the case of verbal WM, alpha suppression in response to the first couple of memory items is inseparable from perception.

A systematic review of the literature identified that alpha during retention interval does not always change in the same direction, with 80% of studies reporting an alpha increase and the rest showing the opposite pattern in similar conditions (Pavlov & Kotchoubey, 2022). The alpha increase during retention has been associated with the suppression of task-irrelevant cortical areas (Jensen & Mazaheri, 2010) that serves the functions of distractor inhibition and disengagement of external attention during retention period (Clayton et al., 2018; Payne & Sekuler, 2014; Roux & Uhlhaas, 2014; Schroeder et al., 2018). Unlike most previous studies, we did not observe an alpha increase during the retention period. It is puzzling because disengagement of visual cortical areas would potentially benefit performance especially in a task with auditory stimulus presentation. Additionally, we would have expected to observe weaker alpha suppression during encoding and retention of later digits in the sequence in the participants employing the strategy to ignore or suppress new digits while keeping only a limited number of memory items, but this was not the case. Possibly, alpha enhancement during retention interval in WM is not always related to inhibition of task-irrelevant information and related cortical areas.

The continuous suppression of alpha over the duration of the task may reflect preparation for encoding of new items because the length of the sequence was not known to the participants. Notably, as shown in Figures 5 and S3, the most apparent changes in alpha activity were observed during encoding and maintenance of the 6^th^ and 10^th^ digits – the digits following the presentation of the possible final items in the sequences. But this explanation is less convincing for the longest 13-digit sequences. Alpha activity and pupil dilation have different temporal dynamics, and sustained suppression of alpha activity is unlikely to be related to an increasing level of arousal, but short surges of arousal – orienting reaction (expectation violation or surprise) at the 6^th^ and 10^th^ digits might contribute to the observed pattern. Accordingly, the effects in the alpha frequency band may be caused by the orienting reaction and the division of attention between memory items during encoding and rehearsal.

### Oscillations and pupil diameter

Frontal midline theta mirrored the dynamics of pupil dilation with the exception that theta power never reached the baseline level, while alpha activity was not associated with pupil size dynamics. Despite the potential connection, the data on the relationship between brain oscillations and pupil dilation are limited (Dippel et al., 2017; Lin et al., 2018). Referenced studies employed the tasks eliciting transient theta changes like the no-go inhibitory control task (Lin et al., 2018), but continuous oscillatory activity visible during maintenance of WM has never been studied before in relation to pupil dilation.

The correlation between theta and pupil temporal trajectories may point to a shared underlying mechanism. LC-NE system has been argued to be related to pupil diameter (Joshi et al., 2016; Murphy et al., 2014). The activation of locus coeruleus (LC) has been associated with firing patterns in anterior cingulate cortex (ACC), suggesting reciprocal connections of LC-NE with cortex (Joshi & Gold, 2022). At the same time, the relationship between ACC and frontal theta has been demonstrated previously by the means of source modelling (Ishii et al., 1999; Meltzer et al., 2007; Onton et al., 2005), intracranial recordings (Raghavachari et al., 2001), and fMRI (Scheeringa et al., 2009). NE delivery from the LC was suggested to amplify gain in fronto-parietal network (Unsworth & Robison, 2017). LC-NE is important for the maintenance of alertness and sustained attention (Chamberlain & Robbins, 2013; Sara, 2009). One may speculate that the ACC or possibly other prefrontal cortical areas are involved in top-down control of attention and may also be responsible for the maintenance of vigilance during the task via activation of LC. Theta power increase and pupil dilation may emerge from top-down modulation of the release of neuromodulators from LC from ACC. But the lack of robust correlation between theta and performance while the significant relationship between pupil size and performance suggests not fully overlapping mechanisms of the phenomena.

Power in the alpha band showed a different response to the cognitive load, compared to pupil size. One theory connects LC-NE system function and alpha suppression (Dahl et al., 2022; Sharon et al., 2021) stating that LC-NE system modulates processing in cortico-thalamic networks (Rodenkirch et al., 2019; Rodenkirch & Wang, 2020), and alpha activity is a product of multiple thalamic and cortical sources (Bollimunta et al., 2011; Crunelli et al., 2018).

Although this intricate link of LC-NE with alpha suppression is theoretically sound (Dahl et al., 2020, 2022), we have found no evidence for it in the current study.

## Conclusions

To summarize, using concurrent pupillometry and EEG during performance of the digit span task with sequences comprising up to 13 digits, we showed that (1) upon reaching cognitive overload, pupil constricts down to baseline level; (2) temporal dynamics of pupil size and frontal midline theta activity are similar, suggesting that they have similar neural mechanisms; (3) theta activity is above baseline level only when there is a sequence with temporal order to memorize (no apparent theta for one memory item), indicating the binding function of theta rhythm; (4) as alpha suppression persists and increases at high levels of WM load exceeding normal WM capacity (10 items or more), alpha cannot be associated with attentional focus or distractor suppression.

## Author contributions statement

Alexandra I. Kosachenko: Investigation, Methodology, Data curation, Visualization, Formal analysis, Writing - original draft, Writing - review & editing.

Dauren Kasanov: Investigation, Methodology, Data curation, Writing - review & editing.

Alexander I. Kotyusov: Investigation, Methodology, Data curation, Writing - review & editing.

Yuri G. Pavlov: Conceptualization, Funding acquisition, Data curation, Formal analysis, Project administration, Supervision, Visualization, Methodology, Writing - original draft, Writing - review & editing.

## Data availability statement

The data are publicly available on OpenNeuro (https://openneuro.org/datasets/ds003838).

## Acknowledgements

We are immensely grateful to Niko Busch and Boris Kotchoubey for providing valuable feedback on earlier versions of the manuscript. The research funding from the Ministry of Science and Higher Education of the Russian Federation (Ural Federal University Program of Development within the Priority-2030 Program) is gratefully acknowledged.

