## Supplementary for "EEG and pupillometric signatures of working memory overload"

### Supplementary results

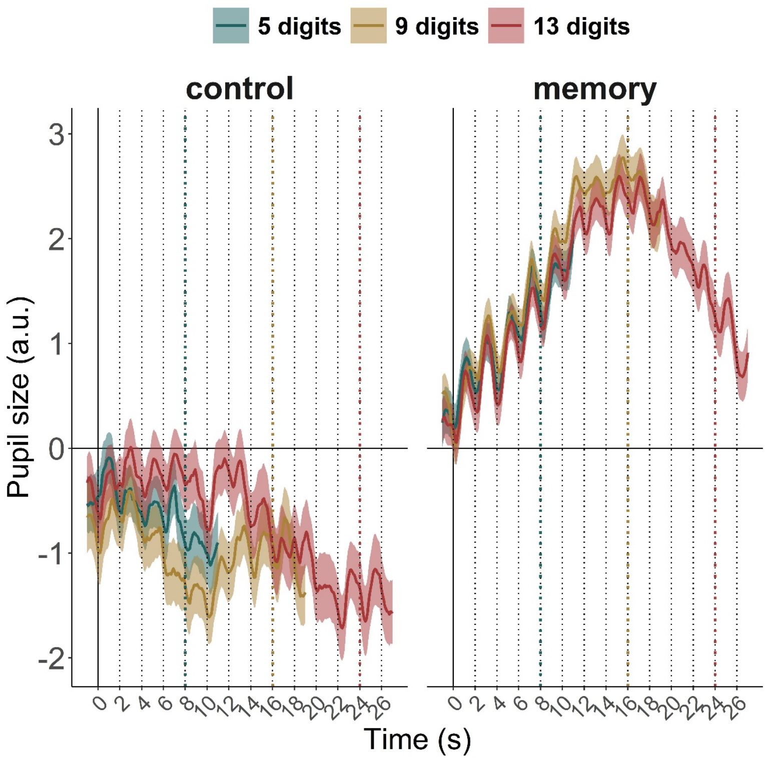


Figure S1 – Pupil size dynamics in the first block of trials. Note that only data from participants with at least two trials per condition was included in the average (N = 61).

### Memory strategy effects

Table S1 – The effect of the strategy to remember only a limited number of the first digits in the sequence or try to remember everything on pupil size

| **Effect** | ***df*** | ***F*** | $\eta_p^2$ | ***p*** |
|---|---|---|---|---|
| Group | 1, 70 | 0.0267 | 0.00038 | 0.8706 |
| Load | 1.8941, 132.58 | 10.8848 | 0.13457 | 5.92E-05 |
| Group x Load | 1.8941, 132.58 | 0.5855 | 0.00829 | 0.5493 |
| Task | 1, 70 | 119.9909 | 0.63156 | < 2.2e-16 |
| Group x Task | 1, 70 | 0.0118 | 0.00017 | 0.9139 |
| Load x Task | 2.3056, 161.39 | 58.0656 | 0.4534 | < 2.2e-16 |
| Group x Load x Task | 2.3056, 161.39 | 0.2654 | 0.00378 | 0.7977 |

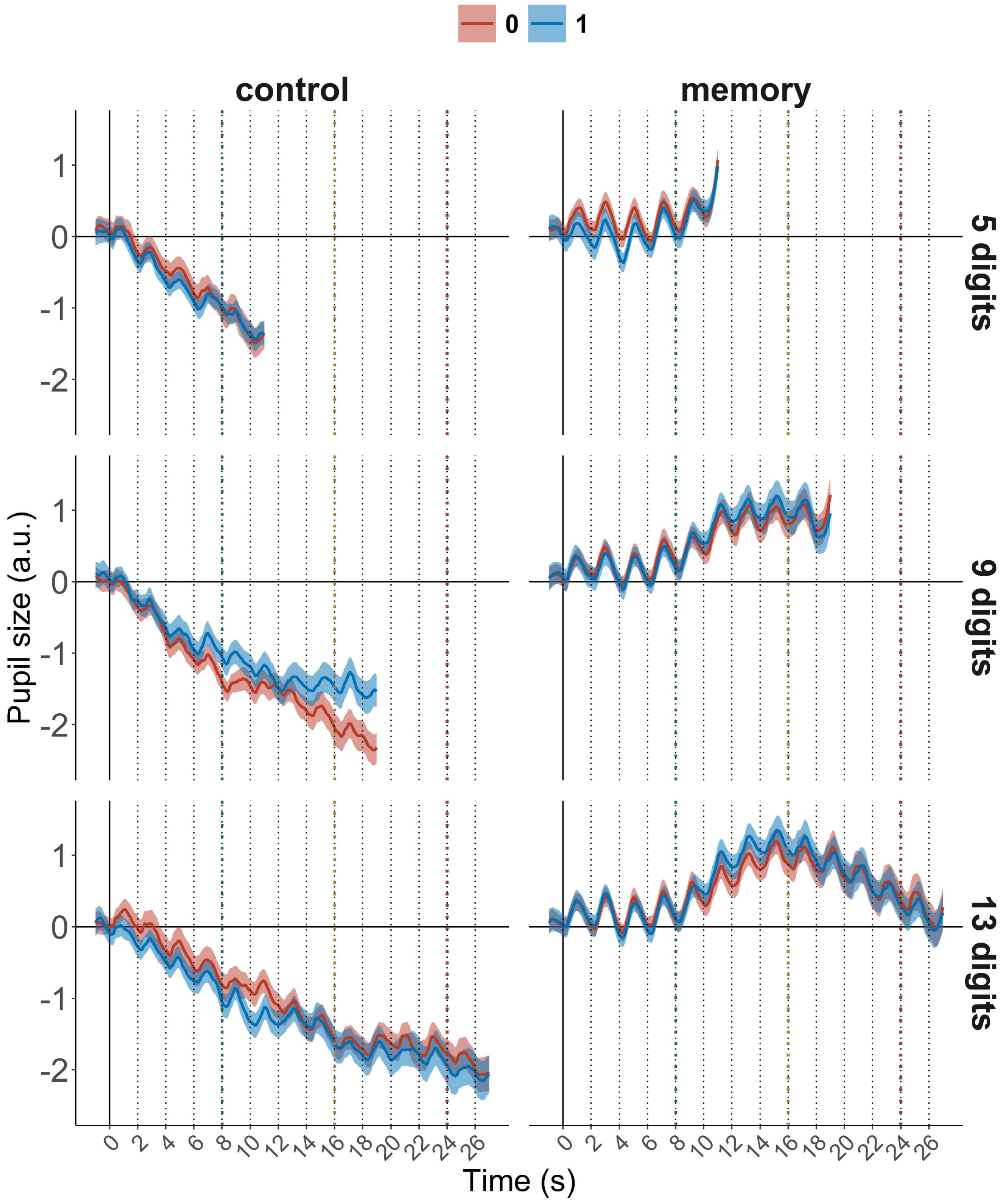


Figure S2 – Group difference strategy to remember only a limited number of the first digits in the sequence (1) or try to remember all presented digits (0)

#### Encoding and maintenance of individual digits

Table S2 – Pairwise comparison of relative theta activity in 2 second time window after digit presentation averaged over all sequences in control condition. Note – uncorrected p-values are reported

| Contrast (digit serial positions) | *t(64)* | *p* |
|---|---|---|
| 1 - 2 | 1.772 | 0.0812 |
| 1 - 3 | 1.196 | 0.2361 |
| 1 - 4 | -0.752 | 0.4551 |
| 1 - 5 | 0.001 | 0.9996 |
| 1 - 6 | -1.136 | 0.26 |
| 1 - 7 | -0.553 | 0.5825 |
| 1 - 8 | -0.512 | 0.6104 |
| 1 - 9 | -1.24 | 0.2196 |
| 1 - 10 | -0.154 | 0.8778 |
| 1 - 11 | 0.7 | 0.4862 |
| 1 - 12 | -0.021 | 0.9829 |
| 1 - 13 | -0.443 | 0.6591 |
| 2 - 3 | -0.298 | 0.7666 |
| 2 - 4 | -2.431 | 0.0179 |
| 2 - 5 | -1.607 | 0.1131 |
| 2 - 6 | -2.38 | 0.0203 |
| 2 - 7 | -1.882 | 0.0644 |
| 2 - 8 | -2.209 | 0.0308 |
| 2 - 9 | -2.755 | 0.0076 |
| 2 - 10 | -1.13 | 0.2628 |
| 2 - 11 | -0.401 | 0.6895 |
| 2 - 12 | -1.078 | 0.2852 |
| 2 - 13 | -1.47 | 0.1464 |
| 3 - 4 | -2.257 | 0.0275 |
| 3 - 5 | -1.316 | 0.1929 |
| 3 - 6 | -2.117 | 0.0382 |
| 3 - 7 | -1.587 | 0.1174 |
| 3 - 8 | -1.456 | 0.1503 |
| 3 - 9 | -1.985 | 0.0515 |
| 3 - 10 | -0.983 | 0.3295 |
| 3 - 11 | -0.22 | 0.8266 |
| 3 - 12 | -0.799 | 0.4274 |
| 3 - 13 | -1.192 | 0.2377 |
| 4 - 5 | 0.91 | 0.3665 |
| 4 - 6 | -0.559 | 0.5782 |
| 4 - 7 | 0.084 | 0.9335 |
| 4 - 8 | 0.154 | 0.8782 |

| | | |
|---|---|---|
| 4 - 9 | -0.618 | 0.5389 |
| 4 - 10 | 0.325 | 0.7465 |
| 4 - 11 | 1.239 | 0.2198 |
| 4 - 12 | 0.449 | 0.655 |
| 4 - 13 | 0.024 | 0.9806 |
| 5 - 6 | -1.263 | 0.2112 |
| 5 - 7 | -0.59 | 0.5574 |
| 5 - 8 | -0.469 | 0.6405 |
| 5 - 9 | -1.274 | 0.2072 |
| 5 - 10 | -0.148 | 0.8828 |
| 5 - 11 | 0.65 | 0.5183 |
| 5 - 12 | -0.022 | 0.9825 |
| 5 - 13 | -0.435 | 0.6647 |
| 6 - 7 | 0.721 | 0.4735 |
| 6 - 8 | 0.745 | 0.4592 |
| 6 - 9 | -0.153 | 0.8785 |
| 6 - 10 | 0.84 | 0.4039 |
| 6 - 11 | 1.851 | 0.0687 |
| 6 - 12 | 0.839 | 0.4046 |
| 6 - 13 | 0.462 | 0.6453 |
| 7 - 8 | 0.098 | 0.922 |
| 7 - 9 | -0.724 | 0.4716 |
| 7 - 10 | 0.295 | 0.769 |
| 7 - 11 | 1.288 | 0.2026 |
| 7 - 12 | 0.409 | 0.684 |
| 7 - 13 | -0.037 | 0.9708 |
| 8 - 9 | -0.993 | 0.3244 |
| 8 - 10 | 0.203 | 0.8399 |
| 8 - 11 | 0.983 | 0.3293 |
| 8 - 12 | 0.315 | 0.7536 |
| 8 - 13 | -0.099 | 0.9215 |
| 9 - 10 | 0.893 | 0.3754 |
| 9 - 11 | 1.625 | 0.109 |
| 9 - 12 | 0.967 | 0.3374 |
| 9 - 13 | 0.571 | 0.57 |
| 10 - 11 | 1.016 | 0.3133 |
| 10 - 12 | 0.126 | 0.8998 |
| 10 - 13 | -0.321 | 0.7493 |
| 11 - 12 | -0.746 | 0.4584 |
| 11 - 13 | -1.301 | 0.1979 |
| 12 - 13 | -0.477 | 0.635 |

Table S3 – Pairwise comparison of relative theta activity in 2 second time window after digit presentation averaged over all sequences in the memory condition. Note – uncorrected p-values are reported

| **Contrast (digit serial positions)** | ***t(64)*** | ***p*** |
|---|---|---|
| 1 - 2 | 1.143 | 0.2575 |
| 1 - 3 | -2.409 | 0.0189 |
| 1 - 4 | -4.518 | <.0001 |
| 1 - 5 | -4.488 | <.0001 |
| 1 - 6 | -6.503 | <.0001 |
| 1 - 7 | -5.895 | <.0001 |
| 1 - 8 | -6.061 | <.0001 |
| 1 - 9 | -5.766 | <.0001 |
| 1 - 10 | -4.293 | 0.0001 |
| 1 - 11 | -4.889 | <.0001 |
| 1 - 12 | -4.136 | 0.0001 |
| 1 - 13 | -3.73 | 0.0004 |
| 2 - 3 | -3.71 | 0.0004 |
| 2 - 4 | -5.92 | <.0001 |
| 2 - 5 | -5.569 | <.0001 |
| 2 - 6 | -7.233 | <.0001 |
| 2 - 7 | -6.602 | <.0001 |
| 2 - 8 | -6.914 | <.0001 |
| 2 - 9 | -6.494 | <.0001 |
| 2 - 10 | -4.69 | <.0001 |
| 2 - 11 | -5.304 | <.0001 |
| 2 - 12 | -4.627 | <.0001 |
| 2 - 13 | -4.143 | 0.0001 |
| 3 - 4 | -3.7 | 0.0004 |
| 3 - 5 | -3.491 | 0.0009 |
| 3 - 6 | -6.035 | <.0001 |
| 3 - 7 | -5.319 | <.0001 |
| 3 - 8 | -5.705 | <.0001 |
| 3 - 9 | -5.601 | <.0001 |
| 3 - 10 | -3.308 | 0.0015 |
| 3 - 11 | -3.325 | 0.0015 |
| 3 - 12 | -2.279 | 0.026 |
| 3 - 13 | -1.973 | 0.0528 |
| 4 - 5 | -0.835 | 0.4071 |
| 4 - 6 | -5.041 | <.0001 |
| 4 - 7 | -4.076 | 0.0001 |
| 4 - 8 | -4.413 | <.0001 |
| 4 - 9 | -3.667 | 0.0005 |
| 4 - 10 | -1.527 | 0.1317 |

| | | |
|---|---|---|
| 4 - 11 | -1.179 | 0.2428 |
| 4 - 12 | 0.173 | 0.8629 |
| 4 - 13 | 0.531 | 0.5972 |
| 5 - 6 | -4.519 | <.0001 |
| 5 - 7 | -4.773 | <.0001 |
| 5 - 8 | -4.861 | <.0001 |
| 5 - 9 | -3.813 | 0.0003 |
| 5 - 10 | -1.324 | 0.1903 |
| 5 - 11 | -0.745 | 0.4592 |
| 5 - 12 | 0.705 | 0.4831 |
| 5 - 13 | 1.002 | 0.3201 |
| 6 - 7 | -0.859 | 0.3934 |
| 6 - 8 | -0.912 | 0.3653 |
| 6 - 9 | 0.082 | 0.9349 |
| 6 - 10 | 0.916 | 0.363 |
| 6 - 11 | 2.015 | 0.0481 |
| 6 - 12 | 3.378 | 0.0012 |
| 6 - 13 | 3.352 | 0.0014 |
| 7 - 8 | 0.026 | 0.9792 |
| 7 - 9 | 0.967 | 0.3374 |
| 7 - 10 | 1.777 | 0.0804 |
| 7 - 11 | 2.751 | 0.0077 |
| 7 - 12 | 4.043 | 0.0001 |
| 7 - 13 | 3.885 | 0.0002 |
| 8 - 9 | 1.043 | 0.301 |
| 8 - 10 | 1.634 | 0.1073 |
| 8 - 11 | 2.814 | 0.0065 |
| 8 - 12 | 4.023 | 0.0002 |
| 8 - 13 | 4.159 | 0.0001 |
| 9 - 10 | 1.032 | 0.3059 |
| 9 - 11 | 2.057 | 0.0438 |
| 9 - 12 | 3.261 | 0.0018 |
| 9 - 13 | 3.802 | 0.0003 |
| 10 - 11 | 0.821 | 0.4147 |
| 10 - 12 | 2.171 | 0.0336 |
| 10 - 13 | 2.566 | 0.0126 |
| 11 - 12 | 2.033 | 0.0463 |
| 11 - 13 | 1.986 | 0.0514 |
| 12 - 13 | 0.502 | 0.6172 |

Table S4 – Pairwise comparison of relative alpha activity in 2 second time window after digit presentation averaged over all sequences in the control condition. Note – uncorrected p-values are reported.

| **Contrast (digit serial positions)** | ***t(64)*** | ***p*** |
|---|---|---|
| 1 - 2 | -1.135 | 0.2606 |
| 1 - 3 | -0.821 | 0.4149 |
| 1 - 4 | -0.031 | 0.9753 |
| 1 - 5 | -0.264 | 0.7924 |
| 1 - 6 | 0.083 | 0.9338 |
| 1 - 7 | -0.855 | 0.3956 |
| 1 - 8 | -0.182 | 0.8559 |
| 1 - 9 | -0.619 | 0.5379 |
| 1 - 10 | 1.452 | 0.1514 |
| 1 - 11 | -0.945 | 0.3483 |
| 1 - 12 | 0.248 | 0.805 |
| 1 - 13 | 0.461 | 0.6465 |
| 2 - 3 | 0.327 | 0.7451 |
| 2 - 4 | 1.086 | 0.2815 |
| 2 - 5 | 0.748 | 0.4573 |
| 2 - 6 | 0.827 | 0.4113 |
| 2 - 7 | -0.376 | 0.708 |
| 2 - 8 | 0.558 | 0.5791 |
| 2 - 9 | -0.084 | 0.9337 |
| 2 - 10 | 2.269 | 0.0266 |
| 2 - 11 | -0.586 | 0.5603 |
| 2 - 12 | 0.775 | 0.4409 |
| 2 - 13 | 1.007 | 0.3177 |
| 3 - 4 | 0.85 | 0.3985 |
| 3 - 5 | 0.491 | 0.625 |
| 3 - 6 | 0.77 | 0.444 |
| 3 - 7 | -0.533 | 0.5956 |
| 3 - 8 | 0.416 | 0.6784 |
| 3 - 9 | -0.247 | 0.8057 |
| 3 - 10 | 2.138 | 0.0363 |
| 3 - 11 | -0.723 | 0.4723 |
| 3 - 12 | 0.695 | 0.4898 |
| 3 - 13 | 0.976 | 0.3327 |
| 4 - 5 | -0.4 | 0.6904 |
| 4 - 6 | 0.124 | 0.9019 |
| 4 - 7 | -1.008 | 0.3173 |
| 4 - 8 | -0.204 | 0.8391 |
| 4 - 9 | -0.749 | 0.4568 |
| 4 - 10 | 1.623 | 0.1095 |

| | | |
|---|---|---|
| 4 - 11 | -1.051 | 0.297 |
| 4 - 12 | 0.297 | 0.7675 |
| 4 - 13 | 0.519 | 0.6055 |
| 5 - 6 | 0.396 | 0.6937 |
| 5 - 7 | -0.901 | 0.371 |
| 5 - 8 | 0.051 | 0.9595 |
| 5 - 9 | -0.566 | 0.5737 |
| 5 - 10 | 1.786 | 0.0789 |
| 5 - 11 | -0.932 | 0.3548 |
| 5 - 12 | 0.475 | 0.6363 |
| 5 - 13 | 0.716 | 0.4768 |
| 6 - 7 | -1.247 | 0.217 |
| 6 - 8 | -0.397 | 0.6925 |
| 6 - 9 | -1.126 | 0.2645 |
| 6 - 10 | 1.66 | 0.1018 |
| 6 - 11 | -1.083 | 0.2827 |
| 6 - 12 | 0.21 | 0.8344 |
| 6 - 13 | 0.51 | 0.6117 |
| 7 - 8 | 1.108 | 0.2721 |
| 7 - 9 | 0.423 | 0.6735 |
| 7 - 10 | 2.326 | 0.0232 |
| 7 - 11 | -0.376 | 0.7078 |
| 7 - 12 | 1.378 | 0.173 |
| 7 - 13 | 1.28 | 0.205 |
| 8 - 9 | -0.832 | 0.4087 |
| 8 - 10 | 1.902 | 0.0617 |
| 8 - 11 | -0.958 | 0.3418 |
| 8 - 12 | 0.427 | 0.6709 |
| 8 - 13 | 0.676 | 0.5014 |
| 9 - 10 | 2.385 | 0.0201 |
| 9 - 11 | -0.641 | 0.5236 |
| 9 - 12 | 0.967 | 0.3369 |
| 9 - 13 | 1.222 | 0.2261 |
| 10 - 11 | -2.599 | 0.0116 |
| 10 - 12 | -1.271 | 0.2082 |
| 10 - 13 | -1.172 | 0.2455 |
| 11 - 12 | 2.139 | 0.0362 |
| 11 - 13 | 1.893 | 0.0628 |
| 12 - 13 | 0.25 | 0.8034 |

Table S5 – Pairwise comparison of relative alpha activity in 2 second time window after digit presentation averaged over all sequences in the memory condition. Note – uncorrected p-values are reported.

| **Contrast (digit serial positions)** | ***t(64)*** | ***p*** |
|---|---|---|
| 1 - 2 | -1.86 | 0.0675 |
| 1 - 3 | -0.956 | 0.3427 |
| 1 - 4 | 0.419 | 0.6763 |
| 1 - 5 | 1.061 | 0.2925 |
| 1 - 6 | 2.664 | 0.0098 |
| 1 - 7 | 3.325 | 0.0015 |
| 1 - 8 | 2.468 | 0.0163 |
| 1 - 9 | 2.641 | 0.0104 |
| 1 - 10 | 4.347 | 0.0001 |
| 1 - 11 | 3.806 | 0.0003 |
| 1 - 12 | 3.518 | 0.0008 |
| 1 - 13 | 3.973 | 0.0002 |
| 2 - 3 | 0.551 | 0.5836 |
| 2 - 4 | 2.011 | 0.0486 |
| 2 - 5 | 2.208 | 0.0308 |
| 2 - 6 | 3.77 | 0.0004 |
| 2 - 7 | 4.643 | <.0001 |
| 2 - 8 | 3.667 | 0.0005 |
| 2 - 9 | 3.805 | 0.0003 |
| 2 - 10 | 5.398 | <.0001 |
| 2 - 11 | 4.92 | <.0001 |
| 2 - 12 | 4.666 | <.0001 |
| 2 - 13 | 4.944 | <.0001 |
| 3 - 4 | 2.356 | 0.0215 |
| 3 - 5 | 3.063 | 0.0032 |
| 3 - 6 | 4.273 | 0.0001 |
| 3 - 7 | 5.012 | <.0001 |
| 3 - 8 | 4.17 | 0.0001 |
| 3 - 9 | 4.209 | 0.0001 |
| 3 - 10 | 5.771 | <.0001 |
| 3 - 11 | 5.186 | <.0001 |
| 3 - 12 | 4.874 | <.0001 |
| 3 - 13 | 5.026 | <.0001 |
| 4 - 5 | 1.145 | 0.2565 |
| 4 - 6 | 3.583 | 0.0007 |
| 4 - 7 | 4.484 | <.0001 |
| 4 - 8 | 3.331 | 0.0014 |
| 4 - 9 | 3.34 | 0.0014 |
| 4 - 10 | 5.228 | <.0001 |

| | | |
|---|---|---|
| 4 - 11 | 4.843 | <.0001 |
| 4 - 12 | 4.407 | <.0001 |
| 4 - 13 | 4.625 | <.0001 |
| 5 - 6 | 2.255 | 0.0276 |
| 5 - 7 | 2.781 | 0.0071 |
| 5 - 8 | 2.374 | 0.0206 |
| 5 - 9 | 2.258 | 0.0274 |
| 5 - 10 | 3.811 | 0.0003 |
| 5 - 11 | 3.17 | 0.0023 |
| 5 - 12 | 2.692 | 0.009 |
| 5 - 13 | 2.973 | 0.0042 |
| 6 - 7 | 1.667 | 0.1004 |
| 6 - 8 | 0.021 | 0.9836 |
| 6 - 9 | 0.075 | 0.9401 |
| 6 - 10 | 3.15 | 0.0025 |
| 6 - 11 | 2.274 | 0.0264 |
| 6 - 12 | 1.241 | 0.2192 |
| 6 - 13 | 1.766 | 0.0821 |
| 7 - 8 | -1.305 | 0.1967 |
| 7 - 9 | -1.133 | 0.2616 |
| 7 - 10 | 2.416 | 0.0185 |
| 7 - 11 | 1.801 | 0.0764 |
| 7 - 12 | 0.656 | 0.5143 |
| 7 - 13 | 1.213 | 0.2297 |
| 8 - 9 | 0.071 | 0.9433 |
| 8 - 10 | 3.189 | 0.0022 |
| 8 - 11 | 2.63 | 0.0107 |
| 8 - 12 | 1.504 | 0.1374 |
| 8 - 13 | 1.888 | 0.0636 |
| 9 - 10 | 3.283 | 0.0017 |
| 9 - 11 | 2.745 | 0.0078 |
| 9 - 12 | 1.485 | 0.1424 |
| 9 - 13 | 1.874 | 0.0655 |
| 10 - 11 | -0.627 | 0.5328 |
| 10 - 12 | -1.488 | 0.1418 |
| 10 - 13 | -0.618 | 0.539 |
| 11 - 12 | -1.34 | 0.1851 |
| 11 - 13 | -0.277 | 0.7826 |
| 12 - 13 | 0.906 | 0.3683 |

Table S6 – Pairwise comparison of pupil size in 2 second time window after digit presentation averaged over all sequences in the control condition. Note – uncorrected p-values are reported.

| **Contrast (digit serial positions)** | ***t(72)*** | ***p*** |
|---|---|---|
| 1 - 2 | 5.964 | <.0001 |
| 1 - 3 | 8.089 | <.0001 |
| 1 - 4 | 9.38 | <.0001 |
| 1 - 5 | 11.528 | <.0001 |
| 1 - 6 | 9.652 | <.0001 |
| 1 - 7 | 9.546 | <.0001 |
| 1 - 8 | 10.159 | <.0001 |
| 1 - 9 | 10.916 | <.0001 |
| 1 - 10 | 9.481 | <.0001 |
| 1 - 11 | 9.112 | <.0001 |
| 1 - 12 | 9.109 | <.0001 |
| 1 - 13 | 9.67 | <.0001 |
| 2 - 3 | 7.755 | <.0001 |
| 2 - 4 | 8.527 | <.0001 |
| 2 - 5 | 10.61 | <.0001 |
| 2 - 6 | 8.838 | <.0001 |
| 2 - 7 | 8.806 | <.0001 |
| 2 - 8 | 9.196 | <.0001 |
| 2 - 9 | 9.929 | <.0001 |
| 2 - 10 | 8.801 | <.0001 |
| 2 - 11 | 8.412 | <.0001 |
| 2 - 12 | 8.521 | <.0001 |
| 2 - 13 | 9.151 | <.0001 |
| 3 - 4 | 5.259 | <.0001 |
| 3 - 5 | 7.825 | <.0001 |
| 3 - 6 | 6.761 | <.0001 |
| 3 - 7 | 7.25 | <.0001 |
| 3 - 8 | 7.833 | <.0001 |
| 3 - 9 | 8.724 | <.0001 |
| 3 - 10 | 7.716 | <.0001 |
| 3 - 11 | 7.333 | <.0001 |
| 3 - 12 | 7.468 | <.0001 |
| 3 - 13 | 8.03 | <.0001 |
| 4 - 5 | 6.384 | <.0001 |
| 4 - 6 | 4.841 | <.0001 |
| 4 - 7 | 5.697 | <.0001 |
| 4 - 8 | 6.61 | <.0001 |
| 4 - 9 | 8.063 | <.0001 |
| 4 - 10 | 7.135 | <.0001 |

| | | |
|---|---|---|
| 4 - 11 | 6.666 | <.0001 |
| 4 - 12 | 6.61 | <.0001 |
| 4 - 13 | 7.307 | <.0001 |
| 5 - 6 | 1.994 | 0.05 |
| 5 - 7 | 3.28 | 0.0016 |
| 5 - 8 | 4.719 | <.0001 |
| 5 - 9 | 6.707 | <.0001 |
| 5 - 10 | 5.375 | <.0001 |
| 5 - 11 | 4.935 | <.0001 |
| 5 - 12 | 4.979 | <.0001 |
| 5 - 13 | 5.839 | <.0001 |
| 6 - 7 | 2.855 | 0.0056 |
| 6 - 8 | 4.468 | <.0001 |
| 6 - 9 | 6.493 | <.0001 |
| 6 - 10 | 4.687 | <.0001 |
| 6 - 11 | 4.287 | 0.0001 |
| 6 - 12 | 4.268 | 0.0001 |
| 6 - 13 | 5.034 | <.0001 |
| 7 - 8 | 3.536 | 0.0007 |
| 7 - 9 | 6.41 | <.0001 |
| 7 - 10 | 3.639 | 0.0005 |
| 7 - 11 | 3.303 | 0.0015 |
| 7 - 12 | 3.427 | 0.001 |
| 7 - 13 | 4.207 | 0.0001 |
| 8 - 9 | 5.121 | <.0001 |
| 8 - 10 | 2.125 | 0.037 |
| 8 - 11 | 1.968 | 0.0529 |
| 8 - 12 | 2.255 | 0.0272 |
| 8 - 13 | 3.208 | 0.002 |
| 9 - 10 | 0.025 | 0.9803 |
| 9 - 11 | 0.105 | 0.9167 |
| 9 - 12 | 0.552 | 0.5829 |
| 9 - 13 | 1.688 | 0.0958 |
| 10 - 11 | 0.169 | 0.866 |
| 10 - 12 | 0.831 | 0.4089 |
| 10 - 13 | 2.353 | 0.0214 |
| 11 - 12 | 1.057 | 0.294 |
| 11 - 13 | 2.565 | 0.0124 |
| 12 - 13 | 2.721 | 0.0081 |

Table S7 – Pairwise comparison of pupil size in 2 second time window after digit presentation averaged over all sequences in the memory condition. Note – uncorrected p-values are reported.

| **Contrast (digit serial positions)** | ***t(72)*** | ***p*** |
|---|---|---|
| 1 - 2 | -0.14 | 0.8893 |
| 1 - 3 | 0.917 | 0.3625 |
| 1 - 4 | -0.31 | 0.7572 |
| 1 - 5 | -1.267 | 0.2093 |
| 1 - 6 | -3.694 | 0.0004 |
| 1 - 7 | -4.734 | <.0001 |
| 1 - 8 | -4.94 | <.0001 |
| 1 - 9 | -4.288 | 0.0001 |
| 1 - 10 | -3.751 | 0.0004 |
| 1 - 11 | -2.758 | 0.0074 |
| 1 - 12 | -1.66 | 0.1012 |
| 1 - 13 | -0.661 | 0.5109 |
| 2 - 3 | 1.819 | 0.0731 |
| 2 - 4 | -0.334 | 0.7392 |
| 2 - 5 | -1.505 | 0.1368 |
| 2 - 6 | -4.156 | 0.0001 |
| 2 - 7 | -5.076 | <.0001 |
| 2 - 8 | -5.108 | <.0001 |
| 2 - 9 | -4.335 | <.0001 |
| 2 - 10 | -3.731 | 0.0004 |
| 2 - 11 | -2.702 | 0.0086 |
| 2 - 12 | -1.617 | 0.1103 |
| 2 - 13 | -0.622 | 0.5359 |
| 3 - 4 | -2.687 | 0.0089 |
| 3 - 5 | -3.553 | 0.0007 |
| 3 - 6 | -6.197 | <.0001 |
| 3 - 7 | -6.794 | <.0001 |
| 3 - 8 | -6.511 | <.0001 |
| 3 - 9 | -5.411 | <.0001 |
| 3 - 10 | -4.661 | <.0001 |
| 3 - 11 | -3.41 | 0.0011 |
| 3 - 12 | -2.205 | 0.0307 |
| 3 - 13 | -1.111 | 0.2702 |
| 4 - 5 | -3.75 | 0.0004 |
| 4 - 6 | -7.255 | <.0001 |
| 4 - 7 | -7.533 | <.0001 |
| 4 - 8 | -6.799 | <.0001 |
| 4 - 9 | -5.431 | <.0001 |
| 4 - 10 | -4.469 | <.0001 |

| | | |
|---|---|---|
| 4 - 11 | -3.048 | 0.0032 |
| 4 - 12 | -1.739 | 0.0863 |
| 4 - 13 | -0.567 | 0.5726 |
| 5 - 6 | -8.31 | <.0001 |
| 5 - 7 | -7.856 | <.0001 |
| 5 - 8 | -6.695 | <.0001 |
| 5 - 9 | -5.064 | <.0001 |
| 5 - 10 | -3.891 | 0.0002 |
| 5 - 11 | -2.443 | 0.017 |
| 5 - 12 | -1.055 | 0.2949 |
| 5 - 13 | 0.158 | 0.8748 |
| 6 - 7 | -5.147 | <.0001 |
| 6 - 8 | -4.299 | 0.0001 |
| 6 - 9 | -2.715 | 0.0083 |
| 6 - 10 | -1.55 | 0.1256 |
| 6 - 11 | -0.143 | 0.8866 |
| 6 - 12 | 1.374 | 0.1736 |
| 6 - 13 | 2.649 | 0.0099 |
| 7 - 8 | -2.368 | 0.0206 |
| 7 - 9 | -0.842 | 0.4028 |
| 7 - 10 | 0.165 | 0.8694 |
| 7 - 11 | 1.545 | 0.1267 |
| 7 - 12 | 3.076 | 0.003 |
| 7 - 13 | 4.296 | 0.0001 |
| 8 - 9 | 0.898 | 0.3722 |
| 8 - 10 | 1.439 | 0.1544 |
| 8 - 11 | 2.92 | 0.0047 |
| 8 - 12 | 4.505 | <.0001 |
| 8 - 13 | 5.825 | <.0001 |
| 9 - 10 | 1.336 | 0.1858 |
| 9 - 11 | 3.315 | 0.0014 |
| 9 - 12 | 5.129 | <.0001 |
| 9 - 13 | 6.525 | <.0001 |
| 10 - 11 | 3.712 | 0.0004 |
| 10 - 12 | 5.606 | <.0001 |
| 10 - 13 | 6.87 | <.0001 |
| 11 - 12 | 4.989 | <.0001 |
| 11 - 13 | 6.781 | <.0001 |
| 12 - 13 | 5.855 | <.0001 |

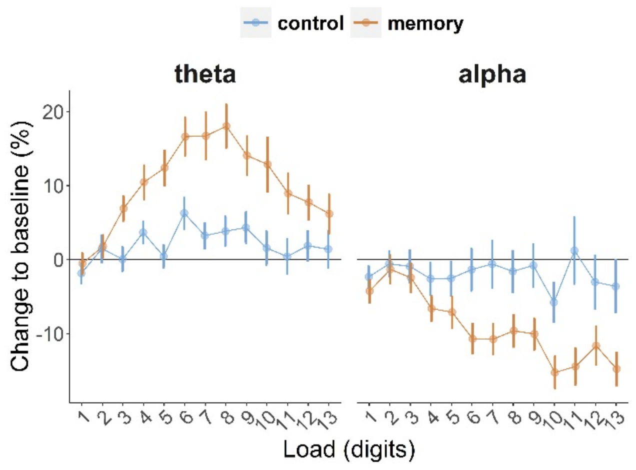


Figure S3 – Average theta and alpha power in 1-2 second time window.

#### Individual differences in physiological activity and memory performance

Table S8 – Internal consistency estimates (Cronbach's alpha) and descriptive statistics of the variables used in the correlational and regression analyses

| variable | mean | SD | skew | kurtosis | reliability |
|---|---|---|---|---|---|
| number of recalled digits in all conditions | 4.12 | 0.81 | 0.2 | -0.12 | 0.95 |
| number of recalled digits in the 5-digit sequences | 4.62 | 0.31 | -1.11 | 1.29 | 0.76 |
| number of recalled digits in the 9-digit sequences | 4.18 | 1.18 | 0.51 | 1.06 | 0.92 |
| number of recalled digits in the 13-digit sequences | 3.57 | 1.13 | 0.25 | -0.79 | 0.92 |
| pupil size in the control task | -1.02 | 1.14 | 0.02 | 1.64 | 0.83 |
| pupil size in the memory task | 0.48 | 1.19 | 0.44 | 0.06 | 0.96 |
| pupil size during retention of 5 digits | 0.36 | 1.47 | 0.3 | -0.24 | 0.92 |
| pupil size during retention of 9 digits | 0.9 | 1.78 | 0.2 | 0.22 | 0.93 |
| pupil size during retention of 13 digits | 0.23 | 1.83 | 0.15 | -0.05 | 0.92 |
| theta power in the control task | 5.02 | 18.98 | 0.69 | 0.56 | 0.41 |
| theta power in the memory task | 13.56 | 14.54 | 0.94 | 0.74 | 0.85 |
| alpha power in the control task | -2.45 | 10.49 | 0.21 | -0.4 | 0.73 |
| alpha power in the memory task | -9.14 | 15.95 | 0.62 | 1.51 | 0.84 |
| alpha power during retention of 5 digits | -12.71 | 22.77 | 0.6 | 0.34 | 0.7 |
| alpha power during retention of 9 digits | -15.49 | 21.85 | 0.55 | -0.21 | 0.65 |
| alpha power during retention of 13 digits | -19.02 | 19.98 | 0.45 | 0.05 | 0.7 |
| theta power during retention of 5 digits | 12.09 | 22.53 | 1.46 | 5.32 | 0.62 |
| theta power during retention of 9 digits | 17.88 | 20.81 | 0.96 | 1.23 | 0.59 |
| theta power during retention of 13 digits | 11.09 | 16.16 | 0.61 | -0.07 | 0.55 |

Table S9 – The comparison low- and high-performance groups (median split) in pupil size

| Effect | *df* | *F* | $\eta_p^2$ | *p* |
|---|---|---|---|---|
| Group | 1, 71 | 0.2525 | 0.00354 | 0.616872 |
| Load | 1.935, 137.39 | 10.8742 | 0.13282 | <0.0001 |
| Group x Load | 1.935, 137.39 | 0.2169 | 0.00305 | 0.798135 |
| Task | 1, 71 | 137.8142 | 0.65998 | < 0.0001 |
| Group x Task | 1, 71 | 7.1175 | 0.09111 | 0.00945 |
| Load x Task | 2.3618, 167.69 | 63.5728 | 0.4724 | < 0.0001 |
| Group x Load x Task | 2.3618, 167.69 | 5.7267 | 0.07464 | 0.002317 |

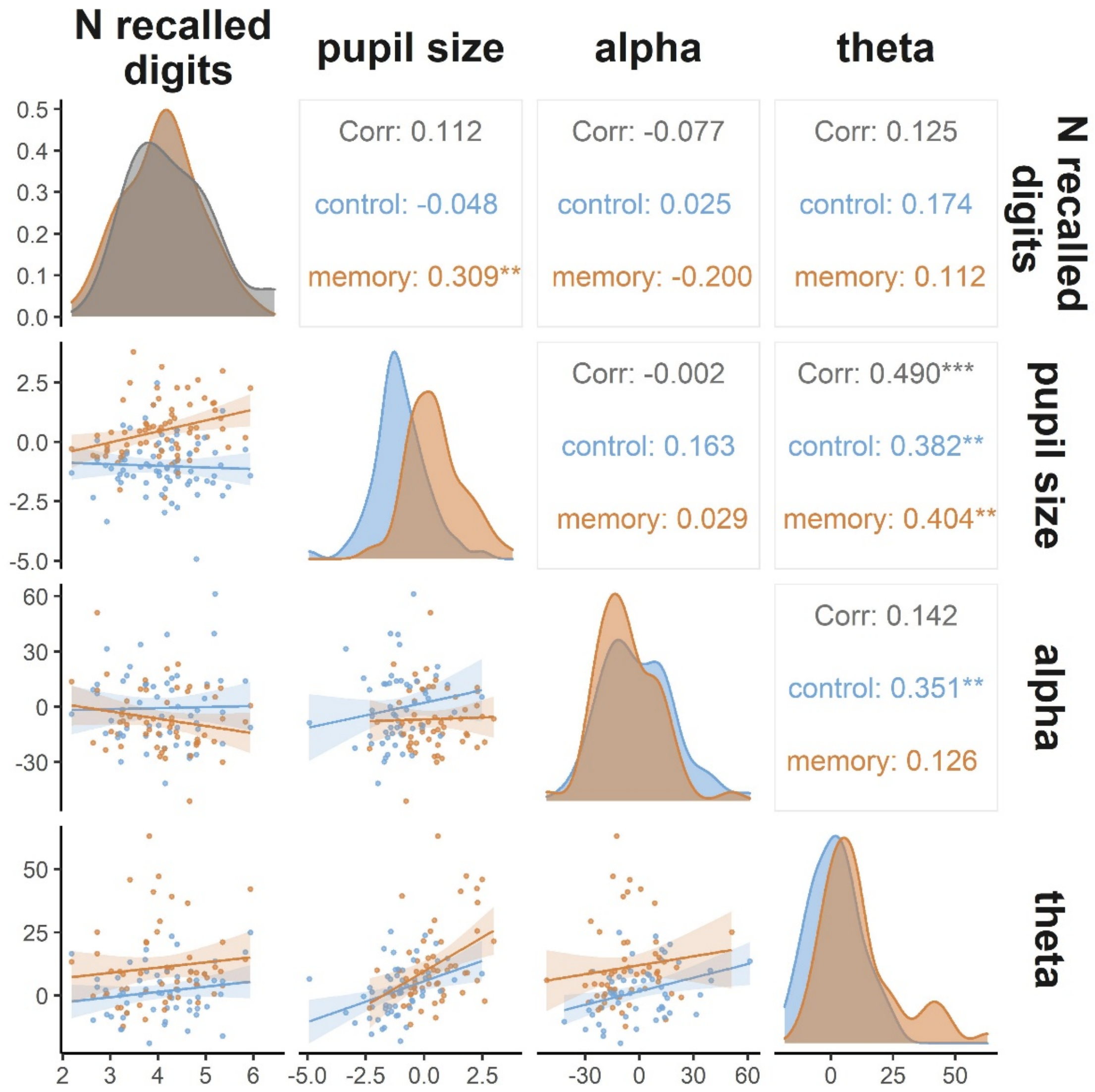


Figure S4 – Correlation of the second second time window for EEG and other variables.

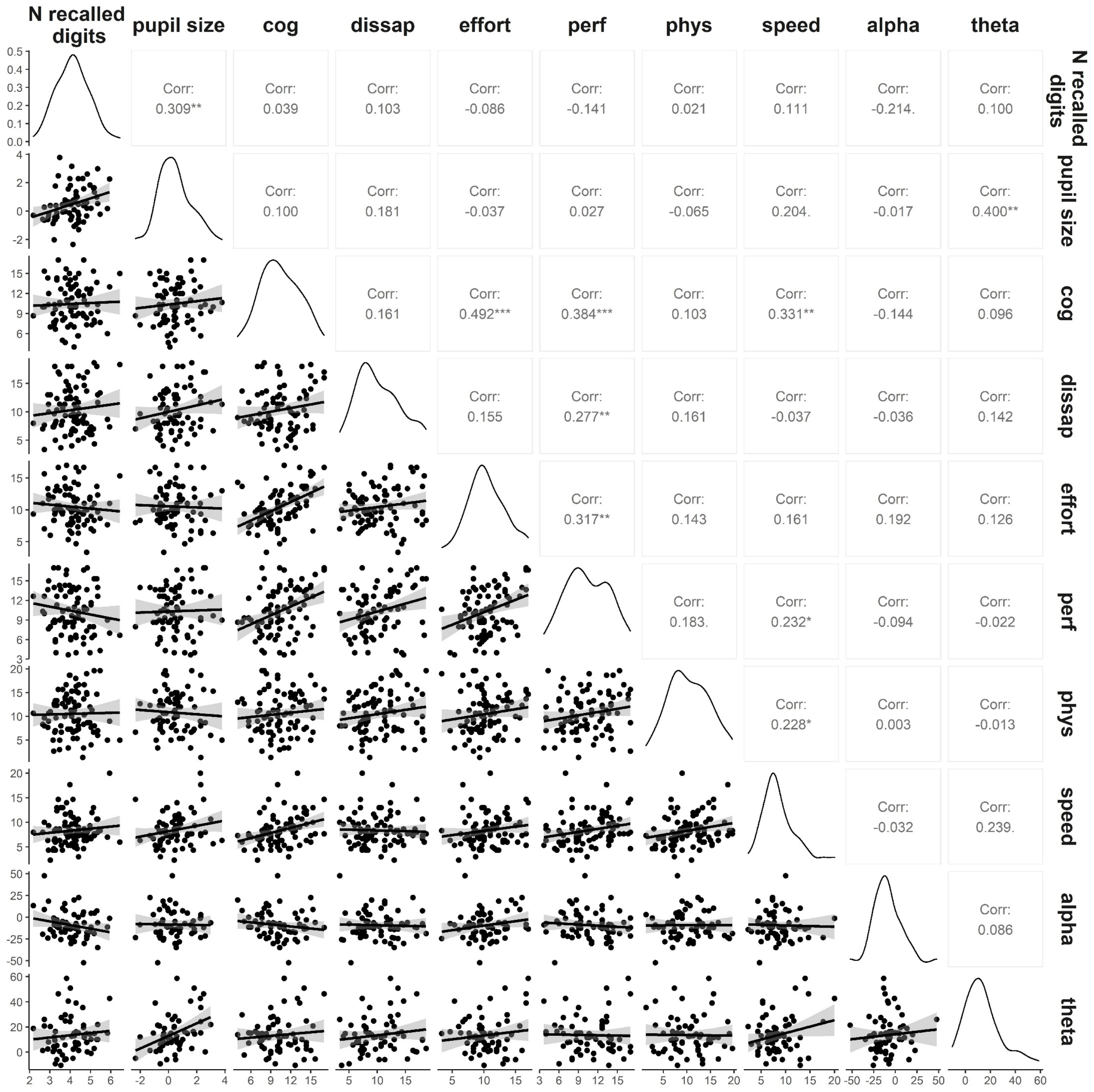


Figure S5 – Pearson correlations of NASA-TLX scales, pupil size, theta and alpha relative spectral power, and number of recalled digits on average across all conditions. NASA-TLX scales: cog – mental load, phys – physical load, speed – temporal demand, perf – perceived overall performance, dissap – frustration.
